# Revisiting the lineages of the Cohanim using data from next-generation sequencing

**DOI:** 10.64898/2025.12.08.692646

**Authors:** J. Lipson, S.G. Bohrer, S. Cohen-Weinstein, L. Cooper, D. Revivo, D. Staetsky, P. Maier, G. Runfeldt, K. Skorecki

**Author notes:** These authors share first authorship (J. Lipson, S.G. Bohrer). **Correspondence:** Joshua Lipson.

## Abstract

**Background:** Early studies using a handful of Y-chromosome markers showed that many self-declared Jewish priests (Cohanim) share related haplotypes, but the limited resolution left open how many priestly lineages exist, when they arose, and how they map onto Jewish population history.

**Objectives:** We used whole-Y sequencing and modern phylogenetic tools to (i) catalogue Y-chromosome clades that are robustly enriched among Cohanim, (ii) date their most-recent common ancestors (TMRCAs) with pedigree-calibrated clocks, (iii) reconstruct their geographic dispersal across the Jewish Diaspora, and (iv) offer a stable, SNP-block nomenclature for future work.

**Methods:** We generated 104 new Big Y-700 sequences from rigorously documented Cohanim spanning 20 diaspora origins, and incorporated 215 legacy samples re-typed for diagnostic SNPs. Phylogenies and branch TMRCAs were built on the FTDNA and YFull backbones. A branch qualified as “Cohen” (CB) if ≥5 men and ≥50 % of branch members self-identified as Cohanim.

**Results:** Nine Cohen branches met these criteria, and for the current study were designated CB-01 to CB-09 with numbering based on antiquity of SNP-block coalescence. CB-01 is the largest (∼45 % of all Cohanim sampled), the most ancient (TMRCA ≈ 850 BCE), and the only lineage present in every major Jewish community. Its internal topology shows explosive branching between 700 and 300 BCE, stasis through the Roman-Byzantine era, and renewed expansion after 800 CE, mirroring known demographic pulses. CB-02 (TMRCA ≈ 700 BCE) and CB-03 (≈ 550 BCE) likewise trace to Iron- or late Classical-period Judea but are geographically narrower. CB-04 – CB-07 are Near-Eastern clades whose priestly status may have crystallized 0–1000 CE, while CB-05, CB-08 and CB-09 appear to derive from Mediterranean host populations. Twenty percent of Cohanim fall outside any current CB, indicating that additional, rarer lineages remain to be characterized.

**Conclusions:** High-resolution sequencing reveals a small set of divergent paternal lines comprising the Jewish priesthood, dominated by a pre-exilic Levantine lineage whose branch structure traces three millennia of geographically dispersed Jewish demographic history. The CB nomenclature, SNP catalogues and open dataset provided here supply a reproducible framework for genetic and historical studies which illustrate the broader power of dense uniparental phylogenies in population-genetic inference.

## Introduction

In one of the earliest applications of phylogenetics used to trace shared uniparental lineages over numerous generations, Y-chromosome sequences and structural variants were examined among Jewish men with a tradition of belonging to the Jewish priesthood, also known as “Cohanim” [1] In this and a landmark follow-up publication, limited sets of phylogenetic markers were able to resolve one large (found among a narrow majority of Cohanim across Jewish communities) and several smaller Y-DNA branches, and enable dating estimates for time to most recent common ancestors (TMRCA) [2]. These branches were found to be significantly enriched among individuals who self-identified as Cohanim, compared to other Jewish and non-Jewish men. Moreover, most members of communities which were otherwise widely dispersed across continents spanning millennia of demographic history, but who identified as Cohanim, clustered into these major and minor branches.

Since the publication of these studies, human population genetics has experienced major advances that include: 1) full genome sequencing accessible and affordable to academic research groups as well as individual consumers [3]; 2) access to very large human genome-wide DNA sequence data sets [4]; 3) techniques for reliable retrieval and analysis of DNA sequence information from small quantities of human skeletal remains [5]; 4) novel data science and bioinformatics tools for analysis of the information resulting from the foregoing advances [6]; [7]. In the case of the Y-chromosome in particular these developments have led to the construction of a staggeringly detailed phylogenetic tree, representing the relationships among publicly accessible human Y-chromosome sequences. To date, the largest such example, hosted by FamilyTreeDNA, comprises 90,000 branches, which together are defined by over 750,000 distinct mutations (**[FamilyTreeDNA May 2025 Report](https://blog.familytreedna.com/ydna-haplotree-90000-branches/)**).

In light of these advances, in the current study we utilized new and existing full Y-chromosome sequences from larger and more diverse data sets to re-examine the detailed phylogenetics of the Cohanim. While this analysis does confirm many of the conclusions of the early studies, it also yields heretofore unappreciated phylogenetic insights of potential historical, cultural and societal interest.

## Methods

### Participants and sampling protocols

Genomic DNA samples were collected with informed consent from male Cohanim, all over the age of 18. All new samples reported in this study were collected in Israel under National Helsinki Committee and Rambam Medical Center Institutional Review Board approval. To ensure the reliability of the priesthood tradition, we only sampled Cohanim who could describe details of their own adherence to practices and rituals of the priestly tradition, and who affirmed similar adherence by both their father and paternal grandfather. Such religious adherence includes avoidance of defilement by contact with the dead, and regular participation in a carefully conducted communal benediction of the congregation during synagogue services. These are practiced most strictly among Cohanim of Orthodox Jewish communities.

We included participants from a wide spectrum of Jewish diaspora geographic origins, with the intention of avoiding disproportionate sampling of Ashkenazim, given their over-representation in previous published studies and in publicly accessible and commercial databases. We also avoided sampling individuals who reported being recently related (that is, within the genealogical timescale).

Participants were supervised in swabbing the inside of their own cheek using a soft disposable swab, and the swab was placed into an Eppendorf tube for genomic DNA extraction and Y-chromosome sequencing. All samples were de-identified, such that the participant data retained by the research team comprised only the paternal geographic origin (**Table 1**). Altogether, samples were collected from 104 previously unsampled individuals. The distribution of reported geographic patrilineal origins (on the basis of paternal grandfather or otherwise earliest known paternal-line ancestor) is as follows: Ashkenazi 43 (41%) with reported patrilines tracing back to Poland, Hungary, Germany, Lithuania, Russia, Belarus, Austria, Ukraine, Slovakia, Romania, Denmark, and the United States; non-Ashkenazi comprising Iran (13), Iraq (11; 3 of whom from Kurdistan), Yemen (8), Tunisia (8; 5 of whom from Djerba), Syria (5), Morocco (4), Afghanistan (3), Algeria (2), Cyprus (1; by self-identification, despite a historical lack of Cypriot Jews in the early modern period), Egypt (1), Greece (1), Israel (1; Musta’arabi Old Yishuv), Lebanon (1), Serbia (1), and Turkey (1). In addition, previously published results of 215 samples of self-reported Cohanim from diverse backgrounds across the Jewish Diaspora (against 1,575 non-priestly Jewish men) which had been collected, reported, and categorized into patrilineal groups [2] on the basis of 75 Single Nucleotide Polymorphisms (SNPs) and 12 Short Tandem Repeats (STRs), were assigned to the more highly resolved Cohen branches, defined according to currently available full Y-chromosome sequence data, and where possible genotyping of branch diagnostic SNPs, as described in Results.

**Table 1:** Summary of Samples.

| Country of Paternal Origin | Number of Samples |
| --- | --- |
| **Ashkenazi** | **43** |
| *Poland* | *14* |
| *Hungary* | *8* |
| *Lithuania* | *5* |
| *Germany* | *5* |
| *Russia* | *4* |
| *Belarus* | *2* |
| *Austria* | *2* |
| *Slovakia* | *1* |
| *Denmark* | *1* |
| *Romania* | *1* |
| **Iran** | **13** |
| **Iraq** | **11** |
| *Kurdistan* | *3* |
| **Yemen** | **8** |
| **Tunisia** | **8** |
| *Djerba* | *5* |
| **Syria** | **5** |
| **Morocco** | **4** |
| **Afghanistan** | **3** |
| **Algeria** | **2** |
| **Cyprus** | **1** |
| **Egypt** | **1** |
| **Greece** | **1** |
| **Israel** | **1** |
| **Lebanon** | **1** |
| **Serbia** | **1** |
| **Turkey** | **1** |

Finally, SNP haplotypes from next-generation Y-chromosome sequences in this study, generated by Gene by Gene as described below, were included in FamilyTreeDNA’s matching databases and publicly viewable phylogenetic trees, allowing the researchers to incorporate information about the ethnogeographic and Cohen status of matching individuals, as well as inferences about the structure of branches as already constituted on FTDNA’s Discover Tree, as described above. In all cases, sample individuals’ branch and terminal SNP assignment were interpreted in the context of their placement within known, publicly available phylogenetic structure.

### Genome Sequencing and variant calling

Sequencing of 104 Y chromosomes was performed at Gene by Gene in Houston, Texas using a combination of Illumina [NovaSeq 6000] and Complete Genomics [DNBSEQ-T7] platforms. We used a target non-recombining region of the Y-chromosome (NRY) region of 15 Mb with a mean 35× sequencing depth, and a minimum of 6× per site. Target-capture sequencing, alignment, and variant calling followed the Big Y-700 protocol described in [8]. The GRCh38 reference genome was used for alignment [9]. For each sample we also generated a total of 700+ Y-STR marker values.

### SNP-based phylogenetic dating

For the Time to Most Recent Common Ancestor (TMRCA) estimates, we followed the method described in [10]. Briefly, we constructed a backbone tree with calibration points for the major clades of the Y-DNA tree using BEAST v2.5.2 [11]. We used a combination of SNP and STR markers to construct Mean Path Lengths (MPLs), which describe the mutational length from each clade to its leaves. The Markov Chain Monte Carlo (MCMC) analyses were run under a strict clock model with a uniform prior on the mutation rate. For SNPs, we applied a mean mutation rate of 0.76 × 10⁻⁹ substitutions per site per year, consistent with pedigree-based estimates, while STRs were modeled with locus-specific mutation rates. We used coalescent constant-size tree priors, ran chains for 100 million generations with sampling every 1,000 steps, and assessed convergence by ensuring effective sample sizes (ESS) >200 for all parameters. SNP-based branch lengths were converted from mutations into time using the equation T = S / (μ × C), where S is the number of SNPs, μ is the Y-DNA mutation rate estimated by [12], and C is the number of bases covered in downstream samples. Union and intersect SNP coverage for the FTBED regions (11.25 Mb of NRY) for each leaf and branch was calculated as described by [13]. STR-based branch lengths were calculated by modeling SNP-based TMRCA as a function of STR genetic distance across the tree. We applied dual SNP/STR-estimates for branches younger than 2,000 years old using a GAM model in the mgcv package v1.8.28 [14]. To account for heterotachy, we applied the relaxed clock method of [15], which uses a least squares criterion to minimize branch length discrepancies. We modeled uncertainty (both in interval between mutations, and mutation rate) as a gamma distribution. Further information regarding TRMCA estimates for Cohen Branches (CBs) are provided in the Supplementary Materials.

For further cross-validation of phylogenies and branch age estimates, we also utilized comparison with the YFull phylogenetic tree (https://www.yfull.com/tree/). Sequence alignment was also cross-validated against ISOGG and YFull reference phylogenies, as was mutation rate estimation [16]. Combining these methodologies in the current study enables examination of consistency across independent datasets thereby providing a robust framework for dating Cohen-associated Y-chromosome branches.

### Markers, Nomenclature, and Matching with Public Databases

The present study defines branches as monophyletic clades marked by unique blocks of shared single nucleotide polymorphisms (SNPs). SNP based clades were used to improve statistical power and phylogenetic resolution compared to approaches which utilize only short tandem repeat (STR) approaches. Accordingly, the current study delineates Cohanim branches in terms of clades defined by a unique block of SNPs shared by all members of a clade, to the exclusion of all non-members. In some cases, such as that of the largest Cohanim clade described in Results, these SNP blocks can comprise dozens of unique, branch-defining SNPs, but have often already been given a shorthand name based on a single SNP within the block. The International Society of Genetic Genealogy (ISOGG) notation begins branch names with an uppercase letter for the crown-level haplogroup (e.g. J), and specifies SNP-defined sub-branches with increasing resolution by adding numbers and letters in an alternating, increasingly lengthy sequence (e.g. J1a2a1a2d2b2b2c2a∼). Consumer population genetic testing and citizen science often combine the name of a crown-level clade defining letter designation (e.g. J), and an alphanumeric name of one of the mutations that defines the branch as distinct from all other branches (e.g. ZS222), into a dash-bound compound name (e.g. J-ZS222) [26] Other third-party organizations, such as the non-profit Jewish genetic genealogy organization Avotaynu, have developed their own forms of notation for naming and cataloging known Jewish Y-DNA branches. For purposes of the current study, a simplified, flexible nomenclature independent of any one convention was adopted. To ensure clarity, consistency, and adaptability for readers of varied backgrounds, Cohanim branches are labeled sequentially as “CB-01,” “CB-02,” and so forth, with CB-01 denoting the earliest Cohen- TMRCA and higher numbers indicating progressively younger branches. It is appreciated that the relative chronological order of TMRCA estimates for these lineages might change, in the wake of future sampling and methodological advances.

We stipulate several conditions which qualify a branch as having a “Cohen” designation for the purposes of this study. First, a CB must be nested within or coextensive with a recognizably Jewish branch, in turn defined as a distinct phylogenetic branch, all or nearly all of whose downstream branches are predominant among Jewish compared to non-Jewish men. We note that the specific aim of this method is to identify distinct “Cohen TMRCAs”, which represent the most remote level under which a secure majority of modern descendants identify as Cohanim - and that in several cases, the Cohen TMRCA used to define a CB is nested downstream of a broader Jewish, non-exclusively Cohen-associated TMRCA. Second, recognizing the practical limitations imposed by sample sizes - especially among non-Ashkenazi Jewish communities where sampled individuals are fewer- a minimum threshold of five identified Cohanim (either from the present study sample, or from corresponding connections in the FTDNA database) was set to define a Cohen Branch (CB). A smaller number, such as one or two individuals, would fail to meaningfully represent a distinct lineage group. Conversely, selecting a significantly larger number would risk excluding authentic branches due to insufficient sampling from smaller communities, thus limiting the inclusivity and representativeness of the analysis. Third, in addition to this minimum requirement, at least 50% of the branch members must self-identify as Cohanim to qualify as a CB. This threshold ensures the distinctiveness and representative validity of each identified branch. We expand on the impact of using higher thresholds “Limitations and Future Directions” section of the Discussion.

SNP haplotypes from whole Y-chromosome sequences in this study were compared with self-reported Cohanim in publicly accessible databases, primarily drawing upon citizen-science projects at FamilyTreeDNA. In a few instances, other databases (e.g. YFull) built on comparable methods were consulted. This comparison enables: 1) assessing congruence between self-reported and investigator-determined Cohanim tradition, 2) identifying additional sub-branches, and 3) exploring broader geographic origins of key branches. This comparison allowed for the enrichment of our sample size, and thus more robust inferences about Cohen status, geographical distribution, and phylogenetic depth of CBs identified in this study.

Finally, we stress that the purpose of this inquiry is strictly *descriptive*: it is not our intention to designate particular Y-DNA branches as representing “authentic" Cohanim in either a historical or ritual sense. For example, a lower CB branch number is not intended to be used to indicate historical, social or ritual preeminence. Rather, population genetics can be informed by and can itself contribute information relevant to population history, while personal identity is an individual matter informed by a variety of sources which include also population and family history and tradition.

## Results

Across a wide sample spanning nearly the entire Jewish Diaspora, we identified nine distinct branches robustly associated with Cohanim status, based on the criteria specified (**Table 2**). These are named and reported in descending order of antiquity based on estimated branch age. **Table 3** shows that some of these branches are shared among geographically and historically distant Jewish communities; others, by contrast, are particular to a single geographical region. **Table 3** also shows SNP haplotype molecular clock-based dating estimates for the time to most recent common ancestor (TMRCA) for all samples in each given CB, which trace back to time periods ranging from the early first millennium BCE for CB-01, the most ancient, to the 16^th^ century CE for the most recent.

**Table 2:**
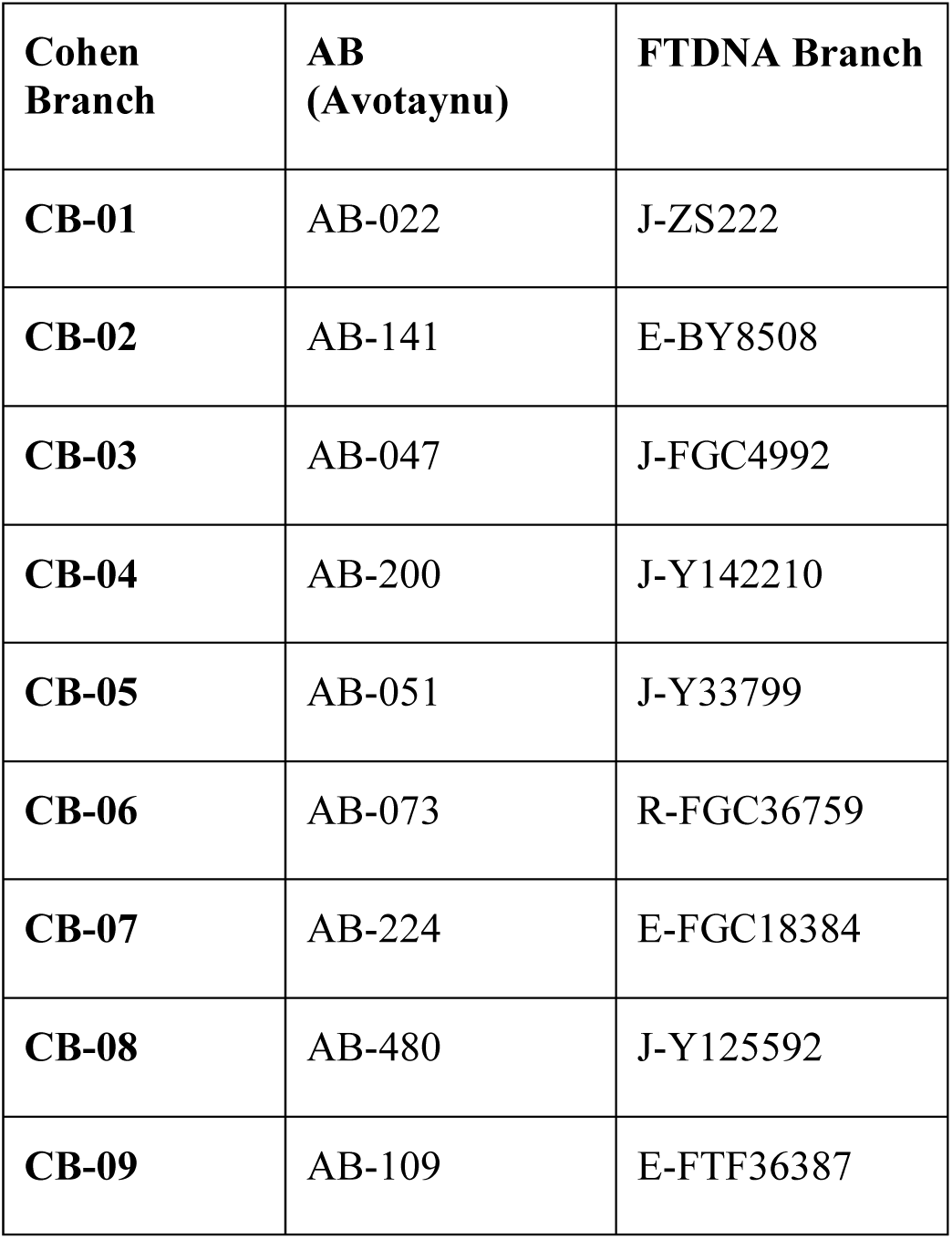
Comparison reference table: CBs, ABs, and FTDNA branches.

| **Cohen Branch** | **AB (Avotaynu)** | **FTDNA Branch** |
| --- | --- | --- |
| **CB-01** | AB-022 | J-ZS222 |
| **CB-02** | AB-141 | E-BY8508 |
| **CB-03** | AB-047 | J-FGC4992 |
| **CB-04** | AB-200 | J-Y142210 |
| **CB-05** | AB-051 | J-Y33799 |
| **CB-06** | AB-073 | R-FGC36759 |
| **CB-07** | AB-224 | E-FGC18384 |
| **CB-08** | AB-480 | J-Y125592 |
| **CB-09** | AB-109 | E-FTF36387 |

**Table 3:** CBs (Cohen branches) defined in the present study.

| **Cohen Branch** | **FTDNA Branch** | **TMRCAs Estimate Oldest (95%)** | **TMRCAs Point Estimate** | **TMRCAs Estimate Youngest (95%)** | **N (testers in present study)** | **% samples in CB for Ashkenazi, non-Ashkenazi, and total samples of current study** | **Communities Represented** | **Likely Origin** |
| --- | --- | --- | --- | --- | --- | --- | --- | --- |
| **CB-01** | J-ZS222 | 1291 BCE | 834 BCE | 440 BCE | 47 | Ashkenazi 64<br>Non-Ashkenazi 33<br>Total 45.19 | All major communities | Israelite/Judean |
| **CB-02** | E-BY8508 | 1548 BCE | 691 BCE | 17 BCE | 4 | Ashkenazi 0<br>Non-Ashkenazi 6.5<br>Total 3.85 | Turkish (Antep),<br>Syrian,<br>Egyptian,<br>Yemenite,<br>Bukharian | Israelite/Judean |
| **CB-03** | J-FGC4992 | 981 BCE | 534 BCE | 152 BCE | 6 | Ashkenazi 14<br>Non-Ashkenazi 0<br>Total 5.77 | Ashkenazi,<br>Moroccan,<br>Greek,<br>Turkish,<br>Mountain,<br>Indian | Israelite/Judean |
| **CB-04** | J-Y142210 | 687 BCE | 1 CE | 537 CE | 5 | Ashkenazi 0<br>Non-Ashkenazi 8.2<br>Total 4.81 | Iraqi, Iranian,<br>Afghan | Near Eastern/<br>Classical Judean |
| **CB-05** | J-Y33799 | 27 BCE | 376 CE | 706 CE | 3 | Ashkenazi 7<br>Non-Ashkenazi 0<br>Total 2.88 | Ashkenazi | Mediterranean<br>European |
| **CB-06** | R-FGC36759 | 333 CE | 755 CE | 1085 CE | 1 | Ashkenazi 2.3<br>Non-Ashkenazi 0<br>Total 1 | Ashkenazi | Near Eastern |
| **CB-07** | E-FGC18384 | 700 CE | 1028 CE | 1285 CE | 9 | Ashkenazi 0<br>Non-Ashkenazi 14.75<br>Total 6.86 | Iraqi, Iranian,<br>Afghan | Near Eastern/<br>Classical Judean |
| **CB-08** | J-Y125592 | 1346 CE | 1552 CE | 1705 CE | 5 | Ashkenazi 0<br>Non-Ashkenazi 8.2<br>Total 4.81 | Tunisian<br>(Djerba) | Mediterranean<br>European |
| **CB-09** | E-FTF36387 | 1295 CE | 1560 CE | 1746 CE | 3 | Ashkenazi 0<br>Non-Ashkenazi 4.9<br>Total 2.88 | Syrian<br>(Aleppo) | Mediterranean<br>European |
| **Others** |  |  |  |  | 21 | Ashkenazi 14<br>Non-Ashkenazi 24.6<br>Total 20.2 | Ashkenazi,<br>Yemenite,<br>Iraqi, Serbian<br>(Sephardic),<br>Iranian,<br>Lebanese,<br>Afghan,<br>Musta'arabi |  |

We also note that a significant minority (20%; *n* = 21) of sampled Cohanim belong to branches not listed in **Table 3**. Some of these (5%; *n* = 5) belong to documented Jewish Y-DNA branches, such as E-BY7500 and R-FT1713, which are generally not associated with priestly status. A larger number (15%; *n* = 16), by contrast, belong to small, poorly-characterized branches which do not meet the scale criteria for CBs in the current data set (e.g. a total of 5 or more members).

For each of the CBs respectively we now provide separate descriptions of results that describe: 1) general phylogenetic context, 2) molecular dating-based age estimates, 3) internal sub-branching structure, 4) distribution across different Jewish communities, and 5) putative origin, drawing upon data obtained from matching in publicly available databases as well as original sampling as detailed above. These results are provided in the text for CB-01, CB-02, CB-03, CB-04, CB-05, CB-06, CB-07, CB-08, and CB-09.

### CB-01

We refer to the first Cohen-associated branch identified in the present study, and associated with the Cohen modal haplotype and J-P58 Cohen cluster of earlier studies, as CB-01. CB-01 is currently defined by the Y-SNP terminal J-ZS222 and its associated block of 25 SNPs (**Supplementary Table 1**), a sub-branch of the broadly Near Eastern haplogroup J-P58, likely associated with early Semitic-speaking populations. We identify J-ZS222 as the relevant phylogenetic level for the purposes of this study, as an overwhelming majority of nearly all of its branches are Jewish individuals with traditions of priestly descent. Using the phylogenetic dating described in Methods, for CB-01, the currently estimated Time to Most Recent Common Ancestor (TMRCA) value is **833 BCE (95% confidence interval (1289 BCE to 440 BCE).**

In this regard, CB-01’s closest upstream connection is to a German Jewish individual with no priestly tradition, displayed on FTDNA’s block tree, and identified with further specificity from a match list. This connection is estimated to date to c. 2150 BCE. Several phylogenetic levels above, currently defined by J-ZS241 (dating to c. 2600 BCE), CB-01 shares a branch with individuals from Bronze and Iron Age Megiddo, in Israel, with a genetically East Mediterranean individual from Imperial-era Rome, and with a range of modern Middle Eastern and European individuals [27], [31] Even more proximally, a recently sampled genome from Bronze Age Sidon, in Lebanon (c. 1700 BCE), shares with CB-01 an ancestor at the level currently defined by J-FGC13863 (dating to c. 2250 BCE [29].

#### Unique features

Corresponding to early reports by Thomas et al. (1998) [20], Hammer et al. (2009) [2] reported the “Cohen modal haplotype” in J-P58—most, but critically not all of which, maps onto CB-01—at a frequency of 46.1% across a large sample of Cohanim, with roughly equivalent frequencies among Ashkenazi and non-Ashkenazi Cohanim. At present, our study’s sample reflects a rate of 45% across an independent sample of Cohanim from across a range of Jewish communities, with rates of 63% among Ashkenazi Cohen samples (somewhat but not significantly higher than Hammer et al.’s (2009) [2] rate of 52%, *perhaps* reflecting stricter sampling criteria in the present study) and 33% among a highly heterogeneous sample of non-Ashkenazi Cohanim, ranging in prevalence from 2 of 13 Iranian Jewish sample individuals to 4 of 4 Moroccan Jewish sample individuals (a rate seemingly corroborated by results from 7 of 8 Moroccan Jewish Cohanim in Hammer et al. (2009 [2]).

To date, CB-01 has been found among Ashkenazi, Moroccan, Algerian, Tunisian, Greek (both Sephardic and Romaniote), Bulgarian, Turkish, Syrian, Iraqi, Iranian, Mountain (i.e. Dagestani), Afghan, and Yemenite Jews, as well as currently non-Jewish individuals of Hispanic/Iberian, Southern Italian, Pontic Greek, Lebanese Christian, Palestinian Christian, Moroccan, Iraqi, Ahwazi, and Yemeni Arab origin but ultimately Jewish patrilineal descent.

We also note that while is typically possible to describe the internal structure of a Jewish Y-DNA branch in prose without sacrificing historically relevant complexity (as for the remaining Cohen branches), CB-01 is a clear exception to this rule. Thus, text-based descriptions of its structure is accompanied by graphical representations (**Figures 1** to **4**).

#### Summary of CB-01’s internal structure

CB-01, as currently defined, splits at its top level (c. 833 BCE) into three sub-branches: **J-FT246135**, **J-Z18271**, and a thus far-unnamed branch defined by a single Brazilian individual of Portuguese origin (**Figure 1**). Notably, the first two of these branches coalesce to nearly the same date, c. 682 BCE and c. 689 BCE, respectively. At c. 682 BCE, J-FT246135 splits into 4 distinct primary-level branches, two of which are found in Iraqi and Syrian Cohanim, with two others represented by non-Jewish individuals (J-Z11314 is found in a Kuwaiti individual and a published sample from a Turkmen individual; a basal individual is of Ahwazi Arab origin from Southwestern Iran) (**Figure 2b)**. In a more extreme cases, J-Z18271 splits c. 689 BCE into 11 known primary descendant branches, as shown in **Figure 2a** and described in the **Supplementary Information**. Nine of these 11 branches exclusively or mostly comprise Jewish individuals from across the Jewish Diaspora (with 8 of 9 associated with Cohen status); the remaining two have only yet been identified in Southern European and Near Eastern populations.

We also find suggestive evidence, based on a supplementary analysis of branching events recorded in FamilyTreeDNA’s phylogenetic tree, that CB-01 experienced significant expansion between 600 and 300 BCE, followed by a long slowdown in branching events, followed by a sharp uptick in expansion from the early medieval period through modern times. We also note that based on FamilyTreeDNA’s dating algorithm, 60 sub-branches of CB-01 attested in contemporary individuals already existed as distinct branches by the beginning of the common era. These dynamics are illustrated visually in **Figures 3** and **4**. This pattern is not observed in the other Cohen branches described below, nor in any other studied Jewish Y-DNA branches. A more extensive, historically-informed interpretation is offered in the Discussion section.

### CB-02

This study designates the second Cohen-associated branch as CB-02. Defined by the terminal Y-SNP E-BY8508 and its associated block of 11 SNPs (**Supplementary Table 2)**, CB-02 is a sub-branch of the broadly Northeast African and Near Eastern haplogroup E-V12. For this study, the E-BY8508 block represents the most relevant phylogenetic level, as the majority of its identified members are Jewish individuals with traditions of priestly descent. For CB-02, the currently estimated TMRCA value is 697 BCE (95% confidence interval (1555 BCE to 21 BCE). To date, CB-02 has been found in Bukharian, Yemenite, Turkish, and Syrian Cohanim, as well as Hispanic individuals with no present tradition of Jewish descent. A diagram of CB-02’s phylogenetic structure and affinities is featured in **Figure 5**.

### CB-03

The third Cohen-associated branch in this study is designated CB-03. Defined by the Y-SNP terminal J-FGC4992 and its associated block of and its associated block of 74 SNPs (**Supplementary Table 3**), CB-03 is a sub-branch of the West Asian haplogroup J-M410 (J2a), initially reported in Hammer et al. (2009) [2]. For this study, the J-FGC4992 block represents the most relevant phylogenetic level, with nearly all members having Jewish backgrounds and traditions of priestly descent. CB-03 is currently estimated to have a TMRCA value of 539 BCE (95% confidence interval (987 BCE to 156 BCE). To date, CB-03 has been found in Ashkenazi, Greek, Turkish, Moroccan, and North Caucasian Cohanim, as well as Indian Jews and Hispanic individuals with no present tradition of priestly descent. A diagram of CB-03’s phylogenetic structure and affinities is featured in **Figure 6**.

### CB-04

CB-04, identified by the terminal SNP J-Y142210 and its associated block of 6 SNPs (**Supplementary Table 4**), is part of the Near Eastern J-P58 haplogroup. Specifically, it represents a second, distinct branch of J-P58, alongside CB-01, which can be associated with Cohen status. Individuals under CB-04 report paternal-line ancestry from Iraq, Iran, and Afghanistan, and carry a tradition of priestly descent; additionally, one individual of Armenian origin clusters closely within Jewish variation. For CB-04, the currently estimated TMRCA value is 1 BCE (95% confidence interval (665 BCE to 521 CE). A diagram of CB-04’s phylogenetic structure and affinities is featured in **Figure 7**.

The remaining CBs we identified, which we name CB-05, CB-06, CB-07, CB-08, and CB-09, cannot be identified with priestly status before the common era. We describe these branches below.

### CB-05

The fifth Cohen-associated branch identified here, designated CB-05, is defined by the Y-SNP terminal J-Y33799 and its associated block of 3 SNPs (**Supplementary Table 5**). As of this report, this is a Cohen branch exclusively found among Ashkenazim; however, a Moroccan Jewish individual with no priestly tradition represents a near outgroup. A sub-branch of the ultimately European haplogroup J-L283 (J2b2), this branch was initially reported as J-M12 in Hammer et al. (2009) [2]. Within this branch, nearly all members identify as Jewish and claim priestly descent. For CB-05, the currently estimated TMRCA value is 375 CE (95% confidence interval (29 BCE to 705 CE). A diagram of CB-05’s phylogenetic structure and affinities is featured in **Figure 8**.

### CB-06

CB-06 is defined by the Y-SNP terminal R-FGC36759 and its associated block of 5 SNPs (**Supplementary Table 6**), a sub-branch of the Indo-European-mediated, but heavily Near Eastern haplogroup R1b-Z2103. As of this report, this is a Cohen branch exclusively found among Ashkenazim. For this study, the R-FGC36759-defined block represents the most relevant phylogenetic level, with nearly all members having Jewish backgrounds and priestly descent traditions. CB-06, is currently characterized by an estimated TMRCA value of 758 CE (95% confidence interval (347 CE to 1079 CE). A diagram of CB-06’s phylogenetic structure and affinities is featured in **Figure 9**.

### CB-07

CB-07 is defined by the Y-SNP terminal E-FGC18384 and its associated block of 12 SNPs (**Supplementary Table 7**), within the Near Eastern haplogroup E-M84. All of its known members are Jewish, with origins in Iraqi, Iranian, and Afghan communities. It also represents the second-largest Cohen branch, after CB-01, among non-Ashkenazi individuals sampled in this project, owing to the present sample’s considerable Iranian and Iraqi Jewish representation. For CB-07, the currently estimated TMRCA value is 1026 CE (95% confidence interval (697 CE to 1284 CE). A diagram of CB-07’s phylogenetic structure and affinities is featured in **Figure 10**.

### CB-08

CB-08 is defined by the Y-SNP terminal J-Y125592 and its associated block of 3 SNPs (**Supplementary Table 8**). Part of the J2a-L70 haplogroup, CB-08 is specific to Jewish individuals from the Tunisian island of Djerba, with Cohen traditions. For CB-08, the currently estimated TMRCA value is 1551 CE (95% confidence interval (1345 CE to 1705 CE). A diagram of CB-07’s phylogenetic structure and affinities is featured in **Figure 11**.

### CB-09

CB-09, defined by the Y-SNP terminal E-FTF36387 and its associated block of 27 SNPs (**Supplementary Table 9**), is a branch within the largely Southeast European haplogroup E-V13. Its members largely identify with one large Cohanic family from Aleppo, Syria. For CB-09, the currently estimated TMRCA value is 1559 CE (95% confidence interval (1293 CE to 1746 CE).

Notably, when phylogenetic date estimates for the age of the aforementioned CBs were cross-referenced with estimates generated by YFull, a striking pattern of cross-validation was observed. Between these two distinct methods, TMRCA estimates were never observed to differ by more than 250 years—and without exception fell within each other’s 95% confidence intervals. Above all, this further demonstrates that robust inferences can be made based on FTDNA-derived TMRCA estimates for the Cohen branches described in this study.

### Comparison of prior STR published lineage designations with CB’s delineated in current study

We also drew upon previously published STR and SNP data from a large sample of Cohanim, for the purposes of comparison with the CBs identified and defined in the present study, using next-generation Y-chromosome sequences. In Table S3 of the supplementary materials accompanying Hammer et al. (2009)’s study of Cohanim [2], Y-DNA results are presented for 215 priestly samples, 93 of which are non-Ashkenazi. For each sample, two types of data are provided: 1) **SNP-based haplogroup designation**: a single SNP marker from a set of 20 identified in the study, and 2) **22 STR markers**. For some individuals from this larger sample, next-generation sequences were generated, and published in Behar et al. (2017) [32]. By combining the STR data from the 2009 study with the current scheme, it was possible to determine the CB assignment for most Cohanim in the table originally provided (**Table 5**). Nonetheless, this determination is not always definitive, as the limited number of STR markers reduces resolution, especially in cases with few close samples or ambiguous results.

**Table 4:** Ashkenazi Sub-Branches of CB-01 (J-ZS222)

| **Sub-Branch Name (FTDNA)** | **FTDNA Branch** | **TMRCA Estimate Oldest (95%)** | **TMRCA Point Estimate** | **TMRCA Estimate Youngest (95%)** | **Non-Ashkenazi Presence** | **Cohen Status** |
| --- | --- | --- | --- | --- | --- | --- |
| J-S12192 | J-S12192 | 408 CE | 648 CE | 852 CE | Peruvian, Mexican, Egyptian Karaite, Moroccan Jew |  |
| **J-S12192 -> J-BY68** | **J-BY68** | 788 CE | 993 CE | 1166 CE | N/A | Yes |
| **J-S12192 -> J-ZS2375** | **J-ZS2375** | 678 CE | 893 CE | 1075 CE | N/A | Yes |
| **J-FT19404** | **J-FT19404** | 385 CE | 875 CE | 1242 CE | Greek Jew, Turkish Non-Jew | Yes |
| **J-ZS2432** | **J-ZS2432** | 703 CE | 961 CE | 1173 CE | N/A | No |
| **J-FT269369** | **J-FT269369** | 1379 CE | 1602 CE | 1761 CE | N/A | Yes |
| J-FGC17481 | J-FGC17481 | 66 CE | 507 CE | 859 CE | Algerian Jew, Tunisian Jew |  |
| **J-FGC17481 -> J-Z18288** | **J-Z18288** | 795 CE | 1128 CE | 1383 CE | N/A | Yes |
| **J-FGC17481-> J-Y142323** | **J-Y142323** | 666 CE | 1097 CE | 1414 CE | N/A | Yes |

**Table 5:**
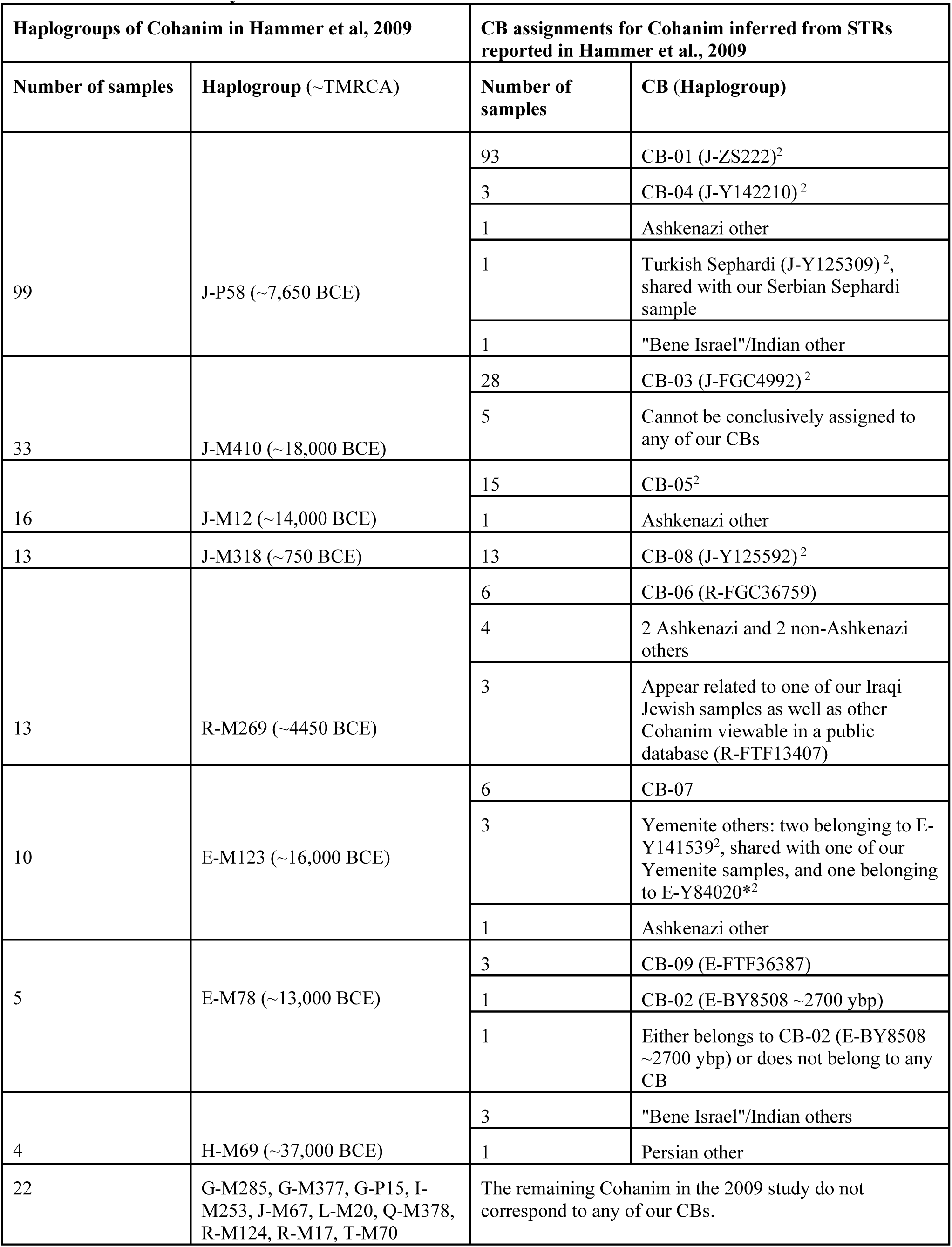
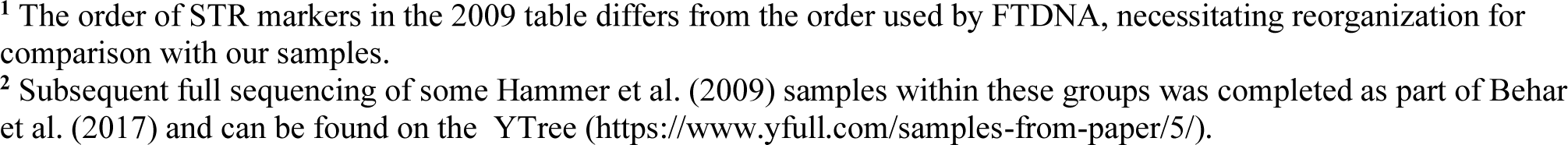
Comparison of prior STR-based lineage designations^1^ (Hammer et al. 2009) with CBs defined in current study.

| Haplogroups of Cohanim in Hammer et al, 2009 |  | CB assignments for Cohanim inferred from STRs reported in Hammer et al., 2009 |  |
| --- | --- | --- | --- |
| Number of samples | Haplogroup (~TMRCA) | Number of samples | CB (Haplogroup) |
| 99 | J-P58 (~7,650 BCE) | 93 | CB-01 (J-ZS222)<sup>2</sup> |
|  |  | 3 | CB-04 (J-Y142210)<sup>2</sup> |
|  |  | 1 | Ashkenazi other |
|  |  | 1 | Turkish Sephardi (J-Y125309)<sup>2</sup>, shared with our Serbian Sephardi sample |
|  |  | 1 | "Bene Israel"/Indian other |
| 33 | J-M410 (~18,000 BCE) | 28 | CB-03 (J-FGC4992)<sup>2</sup> |
|  |  | 5 | Cannot be conclusively assigned to any of our CBs |
| 16 | J-M12 (~14,000 BCE) | 15 | CB-05<sup>2</sup> |
|  |  | 1 | Ashkenazi other |
| 13 | J-M318 (~750 BCE) | 13 | CB-08 (J-Y125592)<sup>2</sup> |
| 13 | R-M269 (~4450 BCE) | 6 | CB-06 (R-FGC36759) |
|  |  | 4 | 2 Ashkenazi and 2 non-Ashkenazi others |
|  |  | 3 | Appear related to one of our Iraqi Jewish samples as well as other Cohanim viewable in a public database (R-FTF13407) |
| 10 | E-M123 (~16,000 BCE) | 6 | CB-07 |
|  |  | 3 | Yemenite others: two belonging to E-Y141539<sup>2</sup>, shared with one of our Yemenite samples, and one belonging to E-Y84020*<sup>2</sup> |
|  |  | 1 | Ashkenazi other |
| 5 | E-M78 (~13,000 BCE) | 3 | CB-09 (E-FTF36387) |
|  |  | 1 | CB-02 (E-BY8508 ~2700 ybp) |
|  |  | 1 | Either belongs to CB-02 (E-BY8508 ~2700 ybp) or does not belong to any CB |
| 4 | H-M69 (~37,000 BCE) | 3 | "Bene Israel"/Indian others |
|  |  | 1 | Persian other |
| 22 | G-M285, G-M377, G-P15, I-M253, J-M67, L-M20, Q-M378, R-M124, R-M17, T-M70 | The remaining Cohanim in the 2009 study do not correspond to any of our CBs. |  |

<sup>1</sup> The order of STR markers in the 2009 table differs from the order used by FTDNA, necessitating reorganization for comparison with our samples.
<sup>2</sup> Subsequent full sequencing of some Hammer et al. (2009) samples within these groups was completed as part of Behar et al. (2017) and can be found on the YTree (https://www.yfull.com/samples-from-paper/5/).

## Discussion

The Cohanim are described in the Hebrew Bible as the priestly class of ancient Israel, traditionally described as being patrilineally descended from biblical Aaron, although critical scholarship has described a potentially more complex situation [33]. In the Pentateuch, the Cohanim are charged with carrying out a variety of ritual and cultic duties, and they recur as key players in descriptions of Israelite society throughout the Hebrew Bible. Documentary evidence from outside the Bible attests to the central place of the Cohanim in the religious and political life of classical Judaea, with priestly elements dominating several early Jewish sects, as well as furnishing the kings of the Hasmonean dynasty (c. 140 to 37 BCE) [34, 35, 36, 37] After the destruction of the Second Temple (70 CE) during the First Roman-Jewish War, the centrality of the Cohanim to Jewish religious life diminished, and Temple rituals gave way to rabbinic and synagogal practices. However, for two millennia throughout the Jewish Diaspora, individuals have continued to identify as Cohanim, a distinction which continues to be of significance in synagogue and lifecycle rituals. At present, it is estimated that between 5 and 10% of Jewish males carry this priestly tradition.

While the early phylogenetic studies cited above yielded results which were consistent with these narratives, it is not at all evident that analyses—based on handfuls of markers and small sample sizes—would yield enough distinctly identifiable extant lineages to robustly capture the complexity of branch histories over these spans of time and space [38]. To this end, the current study utilized full Y-chromosome sequences, allowing more advanced population genetics calculations applied to larger and more diverse data sets to re-examine the detailed phylogenetics of the Cohanim. While this analysis does confirm many of the conclusions of the early studies, it also yielded heretofore unappreciated phylogenetics insights.

The present study offers the first formal update to the state of knowledge on Cohen-associated Y-chromosomal branches since the publication of Hammer et al. (2009) [2], operating with the benefit of next-generation sequencing and phylogenetics, novel sampling, and access to publicly available metadata from consumer and citizen science databases (**[FamilyTreeDNA Discover](https://discover.familytreedna.com/)**; **YFull**). On several points, we find new confirmation for previously-reported findings on the Y-chromosomal landscape of Cohanim, including: 1) the existence of several distinct and unrelated Cohen branches, 2) diversity of Cohen branches *within* individual Jewish communities (e.g. Ashkenazi, Yemenite, and Iraqi Jews), 3) sharing of several Cohen branches *between* different Jewish Diaspora communities, and 4) the unique prominence and spread of one particular Cohen branch, known variously under the labels Cohen modal haplotype, J-ZS222, and in the current paper CB-01. While nine CBs were positively identified using the criteria specified above, this list is not intended to be permanently exhaustive, and is likely to expand as small clusters identified in this study reach a critical sampling threshold.

The present study also offers novel evidence, only possible with next-generation sequencing and advanced phylogenetics, for the continuity of multiple branches’ priestly traditions from the first millennium BCE, as well as for the strong possibility of an ancient Israelite/Jewish affinity of several Cohen branches discussed above—as established by broad phylogenetic methods, ancient DNA, and a novel method focused on intra-Jewish coalescence date estimates in the first millennium BCE. CB-01 can be demonstrated with robust confidence to comprise a monophyletic clade, sub-branches and members of which likely coalesce to a common priestly ancestor in the early first millennium BCE, while CB-02, despite a smaller sample size, appears likely to descend from a common priestly ancestor during the same period—albeit at a lower level of confidence. CB-03’s common Jewish ancestor can securely be dated to the mid-1st millennium BCE, but its early priestly pedigree is somewhat less secure.

As discussed in greater detail below, CB-01 in turn provides the most robust signal of male Jewish migrations across the Diaspora. However, other Cohen branches, such as CB-02 and CB-03, also bridge geographically distant Jewish communities (e.g., in the case of CB-02, Bukharian, Yemenite, Turkish, Syrian, and Egyptian Jews) in ways that are rarely observed in the Jewish Y-DNA landscape. Finally, we offer several hypotheses as to the origins of Cohanim branches whose phylogenetic structures and date estimates do not indicate a particular origin among ancient Israelites/Jews. Based on their Near Eastern origin and the depth of their near-exclusively Jewish phylogenetic structure, CB-04 and CB-07 might well have been present in classical Judaea (c. 2nd-1st century BCE) [30]. CB-06 appears to be broadly Near Eastern in origin based on its phylogenetic affinities, but an ancient Jewish pedigree cannot be established. By contrast, CB-05, CB-08, and CB-09 appear to represent possible introgression from non-Jewish populations in the classical-era Mediterranean Diaspora, based on their phylogenetic affinities and ancient DNA [31].

### CB-01: distinctiveness and historical inferences

We dedicate the remainder of the Discussion to CB-01, given its securely antique association with Cohen status, and its unique phylogenetic structure and ethnogeographical distribution. Drawing upon these defining aspects, we attempt to characterize this largest, most remotely coalescing, yet most widely distributed Cohen branch in historical context. Given that its top-level coalescence date (c. 833 BCE) reaches to the early stages of Jewish history, with documented branching events in nearly every successive century, we posit that an analysis of CB-01 offers valuable data on the dynamics of Jewish population history at every stage. We stress caution in drawing direct correspondences between the phylogeny of CB-01 and known developments in Jewish and ancient Israelite history. At the same time, we attempt to present possible formulations which synthesize these threads in light of critical historiography.

CB-01, though one of many Jewish Y-DNA branches of possible ancient Judean origin, is the Y-chromosomal branch whose modern distribution, antique coalescence date, and broadly Near Eastern affinity most strongly suggests an origin in ancient Israel/Judah. This affiliation is best demonstrated by tracing its descendant branches upward. CB-01’s largest, most widespread sub-branch, J-Z18271, consists of 11 sub-branches, 9 of which have a clear Jewish affinity, coalescing to a common ancestor c. 689 BCE. CB-01’s other named, well-characterized sub-branch, J-FT246135, consists of four sub-branches, two of which have a clear Jewish affinity, coalescing to a common ancestor c. 682 BCE (**Figure 2b**). The most straightforward and parsimonious explanation points to a common origin for both branches, and in turn for their common ancestor c. 833 BCE, in an Iron Age Near Eastern population that contributed ancestrally to all major Jewish Diaspora communities. We posit that the only population that fits this description is that of monarchic period (i.e. First Temple period) Israel and Judah [43]. Moreover, while individuals with CB-01-defining markers have not yet been identified in the ancient DNA literature, individuals from Bronze and Iron Age Megiddo in northern Israel as well as Bronze Age Sidon in Lebanon have been identified as belonging to the branch J-ZS241, which is directly ancestral to CB-01—suggesting deep roots for CB-01 in the Levant [27], [29].

The existence of a priestly branch across nearly all Jewish communities, dating to a common ancestor c. 833 BCE, and without close phylogenetic connections to *non-priestly* Jews, allows for several inferences, some affirming and others refuting prior concepts about early Jewish history and the Jewish priesthood. While a link between CB-01 (or any other Cohen branch) and a common ancestor dating to the biblically reckoned time of Aaron cannot be ruled out by genetic data, the present study offers no affirmative evidence of such an identification. A clearer sense of the earliest date of CB-01’s priestly tradition and Bronze Age origins might be determined through future testing, if individuals splitting the block of 25 SNPs (see **Figure 12**) currently comprising CB-01 and widely designated as J-ZS222 (i.e. testing positive for some relevant SNPs, but negative for others) can be identified. The list of these SNPs using their assigned names in the genetic genealogy literature along with their allelic variants is shown in **Supplementary Table 1**.

Traditional Jewish understanding of the biblical narrative identifies the Cohanim as a subset of the Tribe of Levi, one of 12 Israelite tribes, each descended from a different (eponymous) son of the patriarch Jacob (c. mid-2nd millennium BCE). However, CB-01, which can be identified from its top level with priestly affiliation, is not closely connected to any Jewish non-Cohen branches—that is to say, it does not represent a priestly subgroup within wider Jewish variation. This result, taken together with the fact that modern Jews carry a large range of distinct Middle Eastern-origin Y-DNA branches often not phylogenetically related within the last several thousand years (and belonging to different haplogroups entirely), indicates that the patrilineal origins of ancient Israelites were likely genetically heterogeneous.

On the other hand, the present study’s characterization of CB-01 lends strong circumstantial support to a traditional account of the history of the Jewish priesthood from at the latest c. 833 BCE forward, as well as the authenticity of traditions of priestly descent in communities across the Jewish Diaspora. During the well-documented 9th century BCE, the Israelite ancestors of Jews lived in the larger, prosperous northern kingdom of Israel, and the smaller southern kingdom of Judah, centered around Jerusalem and the First Temple. The earliest common ancestor of CB-01 might trace to this setting, among the priestly class of one of these two kingdoms. It is also possible that CB-02 represents a priestly family of different origin from the same period, following historical-critical scholarship that argues for the existence of competing priestly groups in early Israel [39], [40].

By the turn of the 7^th^ century BCE, after the destruction of the kingdom of Israel by the Assyrians, the kingdom of Judah underwent a period of concentrated population growth, as well as multiple centralizing religious reforms, described in the Hebrew Bible and corroborated by historical-critical scholarship [41, 42, 43]. In a possible parallel, CB-01’s two primary branches, J-Z18271 and J-FT246135, are both estimated to coalesce to the first half of the 7th century BCE, as does CB-02 in its entirety. However, by this early date, CB-01 had already split into at least 16 distinct branches (as attested among contemporary individuals), as compared to only 2 for CB-02. It is possible that as early as the latter days of the First Temple period, CB-01 may already have been the dominant priestly branch in Judah. However, this may also be the result of survivorship bias, due to CB-01 sub-branches’ success in later periods.

The 6th and 5th centuries BCE marked another watershed in the history of ancient Israel, its religious landscape, and the role of Cohanim in society, which may be paralleled in the phylogeny of CB-01. With the Babylonian conquest of Judah and destruction of Jerusalem and its Temple in 586 BCE, Jewish political and religious life was radically transformed—with the Davidic royal elite of the monarchic period dethroned in favor of a new civic-religious priestly elite [44]. Six of J-Z18271’s 11 primary branches have estimated top-level coalescence dates during this period, meaning that they began to split into currently-surviving branches at this time. Indeed, based on FTDNA phylogenetic dating estimates, 42 currently extant branches of CB-01 can be said to have already come into existence by the end of the 5th century BCE.

On the eve of the common era, a minimum of 60 distinct branches of CB-01 had already come into existence, as opposed to 3 each for CB-02, CB-03, and CB-04—which may reflect CB-01’s hegemony among the Judean priestly class by the late Second Temple period (**Figures 3** and **4**). However, the branching growth of CB-01 slowed around the beginning of the common era, possibly corresponding to the period that began with the destruction of the Second Temple by the Romans during the First Roman-Jewish War. In the period that followed, Judaism was gradually reshaped around the rabbinic system *in lieu* of the observance of Temple rituals, relegating Cohanim to relatively minor roles in synagogue ritual [45].

Finally, while CB-01’s pace of branching did not tick upward again until the expansion of Jewish Diaspora communities (e.g. in Germany and Iberia) in the early medieval period, its survival in nearly all major Jewish diaspora populations—often in the form of multiple distinct branches per community—attests to its significance as tracer dye for Jewish Diaspora migrations across several eras, as well as of the phylogenetic relationships between different communities [46]. For example, 7 distinct branches of CB-01 have been identified among modern Ashkenazi Jews, some of them sharing no common ancestor more recent than c. 689 BCE (**Table 4**). While beyond the scope of this study, similar lists of CB-01 sub-branches and their close connections can be made for various Jewish Diaspora communities, including Moroccan, Yemenite, Iraqi, and Eastern Sephardic (Greek and Turkish) Jews.

It is noteworthy that CB-01’s low-level presence in a wide range of populations across the Middle East and Mediterranean functions as perhaps the most robust uniparental signal of Jewish ancestry in groups as wide-ranging as Southern Italians, Yemeni Muslims, and Pontic Greeks. Moreover, a sub-branch of CB-01 has been found *in situ* among Levantine Christians of Palestinian and Lebanese descent, which may represent the signature of an antique Jewish component in these populations.

The preservation of many dozen anciently distinct branches of CB-01 among modern populations is unparalleled, surpassing all other Jewish branches by an order of magnitude in this respect. Several possible factors can be proposed to explain this surprising observation. First of all, it stands to reason based on an analysis of branching events that CB-01 had already greatly diversified within Judaea before the migration of most Jews’ ancestors into the Diaspora, possibly owing to elite status in Second Temple Judaean society. Therefore, it is likely that the Judaean base of most Diaspora communities was heavily seeded with CB-01 sub-branches from the start.

Although the historical-genetic synthesis we present above is hypothetical and subject to change, we emphasize both the *sui generis* nature of CB-01, and its unique ability—through high-resolution phylogenetic structure and dating—to shed light on trends in Jewish population history from the monarchic period to the medieval period.

### Conclusion: wherefore diversity?

Two primary facts concern this discussion: the remarkable case of CB-01, and the diversity of Cohen branches across Jewish Diaspora populations. Several lines of evidence demonstrate CB-01’s unique status among Cohen branches. First, the present study sample, as well as public available data, prove it to be the largest branch among both Ashkenazi and non-Ashkenazi Cohanim. Second, it is found among a wider range of Jewish Diaspora populations than any other Jewish Y-DNA branch, including all other Cohen branches. Third and finally, based on advanced phylogenetic dating algorithms, CB-01 appears to have the earliest coalescence date of any Cohen branch—one which may correspond to the period of the First Temple priesthood—and shows extensive signs of ancient diversification.

However, the results of our investigation also require us to ask the question: in light of CB-01’s prominence, what explains the overall diversity of distinct Y-DNA branches in our sample of Cohanim? Over half of sampled Cohanim are divided across CB-02 through CB-08, as well as smaller branches that do not meet our scale criteria, and also several Jewish Y-DNA branches *not* typically associated with Cohen status.

First, with at least two Cohen branches (CB-01 and CB-02) appearing to trace their priestly pedigree back to early 1^st^ millennium BCE priestly ancestors, we note that a wide range of historians and Bible scholars have suggested the possibility of multiple, unrelated priestly groups in early Israel. And second, while Jewish law precludes the possibility of acquiring priestly status by any means other than descent, it is possible that at various points in Jewish history, priestly status was either adopted out of necessity in the absence of genealogic Cohanim (such as is told in modern Persian Jewish lore, which might explain the preponderance of CB-04 and CB-07 among Iranian and Afghan Jews, as well as in the case of the extinction of the original Samaritan Cohen line in the 17th century), or fabricated by individuals for reasons that have been lost to history. Additionally, Y-DNA branches might have become associated with Cohen status as a result of sporadic NPEs (non-paternity events)—whether ultimately of ancient Judaean origin, or incorporated into Jewish communities through intermarriage and conversion in the Greco-Roman and Babylonian Diasporas.

### Future directions and disclaimer

The current study offers a system of stable, parsimonious nomenclature for Y-DNA branches robustly associated with Cohen status, that facilitates comprehension of the study results for readers without extensive genetic genealogy experience. We are hopeful that this will encourage researchers from many disciplines to pursue studies that will further advance resolution of branching patterns and aspects of demographic history. We also acknowledge that future research studies aimed at characterizing the genetic landscape and origins of Cohanim or Jews at large may or may not choose to continue to use this nomenclature scheme. This scheme is not intended to establish a fixed, invariant set of lineages with Cohen status, but rather, to represent the state of knowledge about the Y-chromosomes of Jews who identify as Cohanim, as of 2025, and to provide a foundation for future inquiries.

A potential limitation of the current study relates to the choice of sample number and percentage thresholds used to establish Cohen lineage and branch designations. As outlined in Methods for purposes of the current study, we used the value of 50% identification as Cohanim for a branch to be designated as a Cohen branch. While 50% was chosen by virtue of signifying a simple majority, applying higher thresholds did not exclude any of the larger groups which met the 50% threshold, up to and through for thresholds of 75-80%. Since these estimates are based on aggregate public databases with variable reporting of genealogical status, we considered that thresholds exceeding 80% would risk excluding valuable information, since attrition of knowledge of Cohanim ancestral affiliation of 20% during the passage of generational time is not surprising and consistent with identifiable lineage attrition predictions [36]. We acknowledge the inherent trade-offs in choosing the cutoff of five individuals. If the threshold were increased significantly, many legitimate but smaller branches, particularly in undersampled non-Ashkenazi communities, would be excluded. Conversely, lowering this threshold would risk erroneously defining branches based on too few individuals, thereby compromising robustness and potentially introducing noise or individual anomalies into the analysis. Consequently, the selection of five individuals strikes a balance between inclusivity for minority populations and methodological rigor, making it the most logical compromise given current sampling constraints. We expect that expansion of sample numbers in the future will allow detailed sensitivity analysis for these thresholds. Additionally, with increased sampling, it should be possible to characterize additional Cohen branches, to clarify the phylogenetic boundaries of Cohen status within wider Jewish branches (e.g. CB-03), and to shore up levels of confidence in the ascription of Cohen status to certain ancient phylogenetic levels (as in CB-02).

Importantly, coalescence timing resolution of CB-01 (as well as other multi-SNP designated CBs) will be advanced by identification of individuals who “split” the phylogenetic SNP blocks that define the Cohen branches discussed above—with a special focus on CB-01, aimed at identifying the maximum antiquity of its Jewish and priestly status. Future studies may attempt to identify such individuals in extant databases using a novel protocol of multiplex PCR reactions which read out the allele status at all relevant block-relevant SNPs, without the necessity for whole Y-sequencing. Moreover, the publication of additional Y-chromosome sequences from remains of ancient and medieval individuals from Israel and the Jewish Diaspora may offer direct evidence for the historical background of documented Cohen branches.

Finally, we do not intend for the conclusions reported in this study to be used to confer special social or ritual status on individuals belonging to one Cohen branch or another. It is widely established that patrilineal *tradition* is the determinant of ritual Cohen status in the Jewish religion; the overlap between tradition, genetic markers, historical inference, and ritual status is likely to be partial at best. While data generated and presented here can be used by interested scholars in many disciplines, we emphasize that we do not intend for “CB-01” or any other designation to confer special status. Rather, the choice of how to make sense of these results in the context of modern life is a matter of personal choice for private individuals and their respective communities.

## Supporting information

Supplementary Informaton

## Conflict of Interest

The authors declare that the research was conducted in the absence of any commercial or financial relationships that could be construed as a potential conflict of interest.

## Author Contributions

JL and SGB are recognized as co-first authors for equal levels of contribution. JL: Original draft, review & editing, conceptualization, formal analysis, investigation, methodology, visualization. SB: Review & editing, conceptualization, data curation, formal analysis, investigation, methodology, project administration, resources, validation. SC-W: Review & editing, investigation, resources. LC: Review & editing, formal analysis, visualization. DR: Review & editing, formal analysis. DS: Review & editing, conceptualization, methodology. PM: Review & editing, conceptualization, data curation, investigation, methodology, resources, software, validation. GR: Review & editing, conceptualization, data curation, investigation, methodology, resources, software, validation. KS: Review & editing, conceptualization, data curation, formal analysis, funding acquisition, investigation, methodology, project administration, resources, supervision.

## Funding

The present project was made possible by funding by the research authority of Bar-Ilan University, including a post-doctoral fellowship award to co-first author SGB.

## Acknowledgments

The authors want to offer acknowledgement to Suzi Nicolau, for her administrative assistance, and Revital Shemer, for her technical assistance and input.

## Supplementary Material

Supplementary information and tables are included as separate files.

## Data Availability Statement

Sequencing results, including Y-STR profiles and initial phylogenetic placement on the FamilyTreeDNA (FTDNA) Discover tree, were uploaded to a dedicated project website (https://www.familytreedna.com/groups/dnaprojects-2024/about), which will become publicly accessible upon publication of this study.

## Notes

### Competing Interest Statement

The authors have declared no competing interest.

