## Supplementary material for "Revisiting the lineages of the Cohanim using data from next-generation sequencing": Figures

### Slide 1
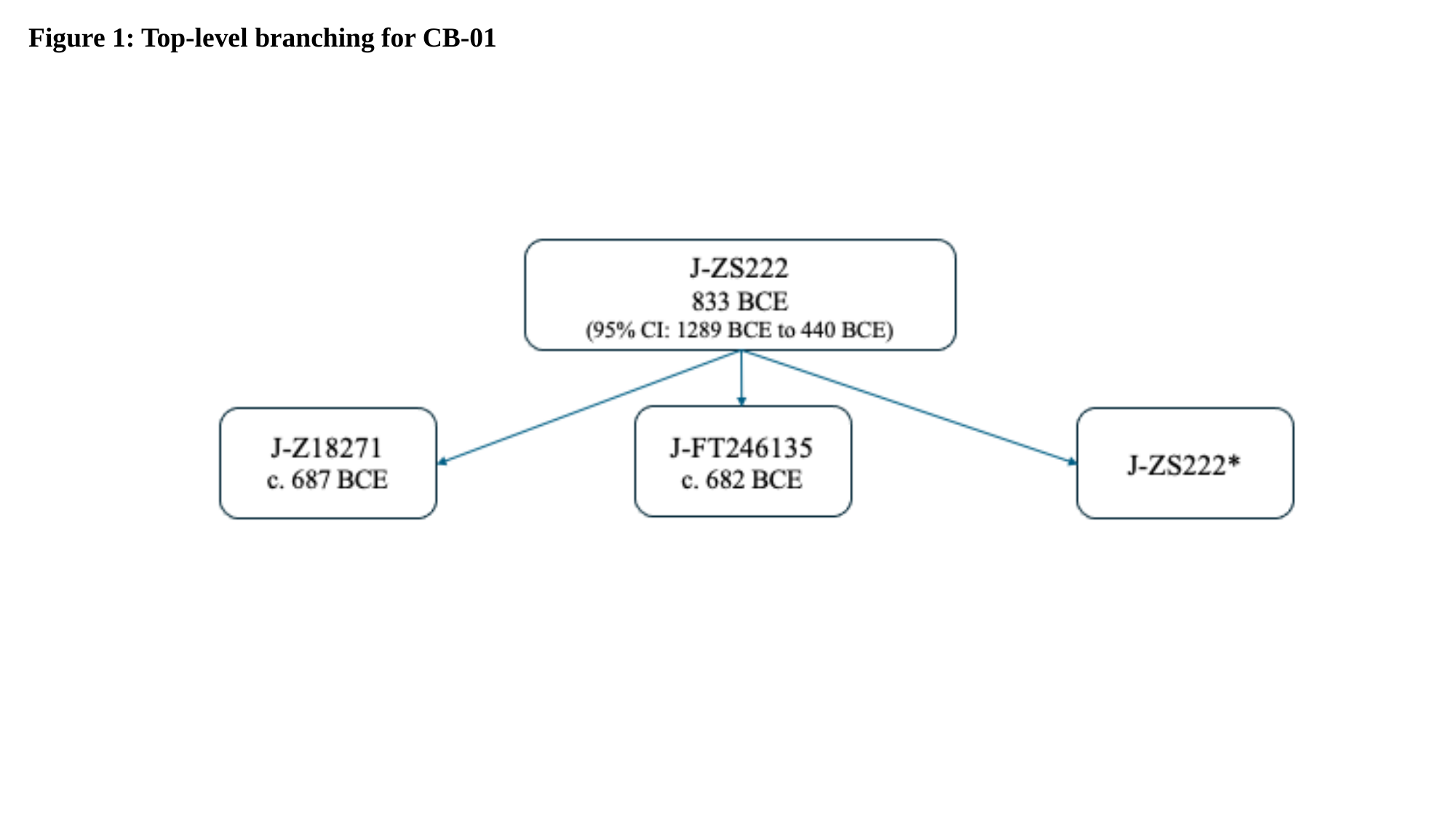

Figure 1: Top-level branching for CB-01

### Slide 2
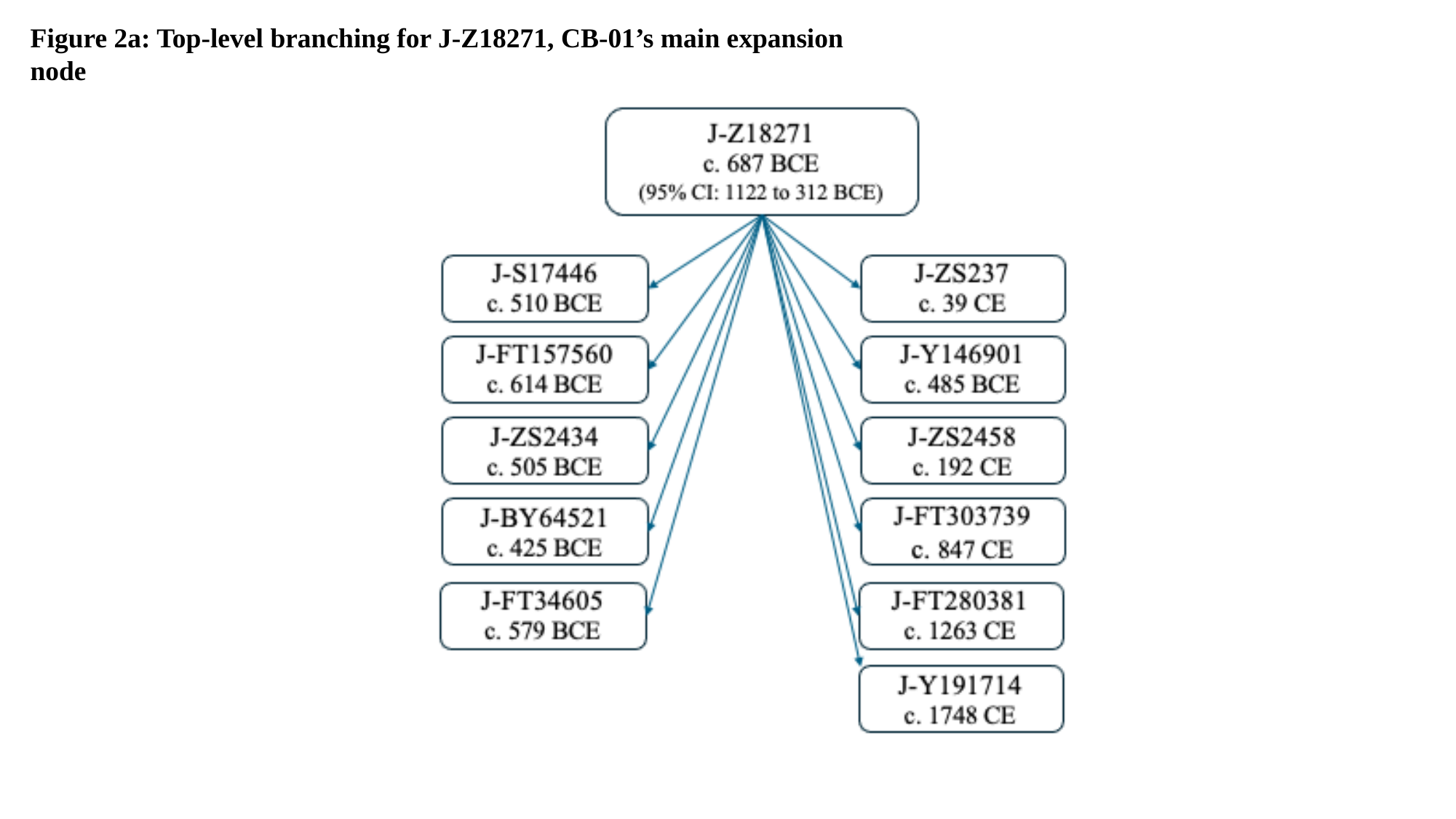

Figure 2a: Top-level branching for J-Z18271, CB-01’s main expansion node

### Slide 3
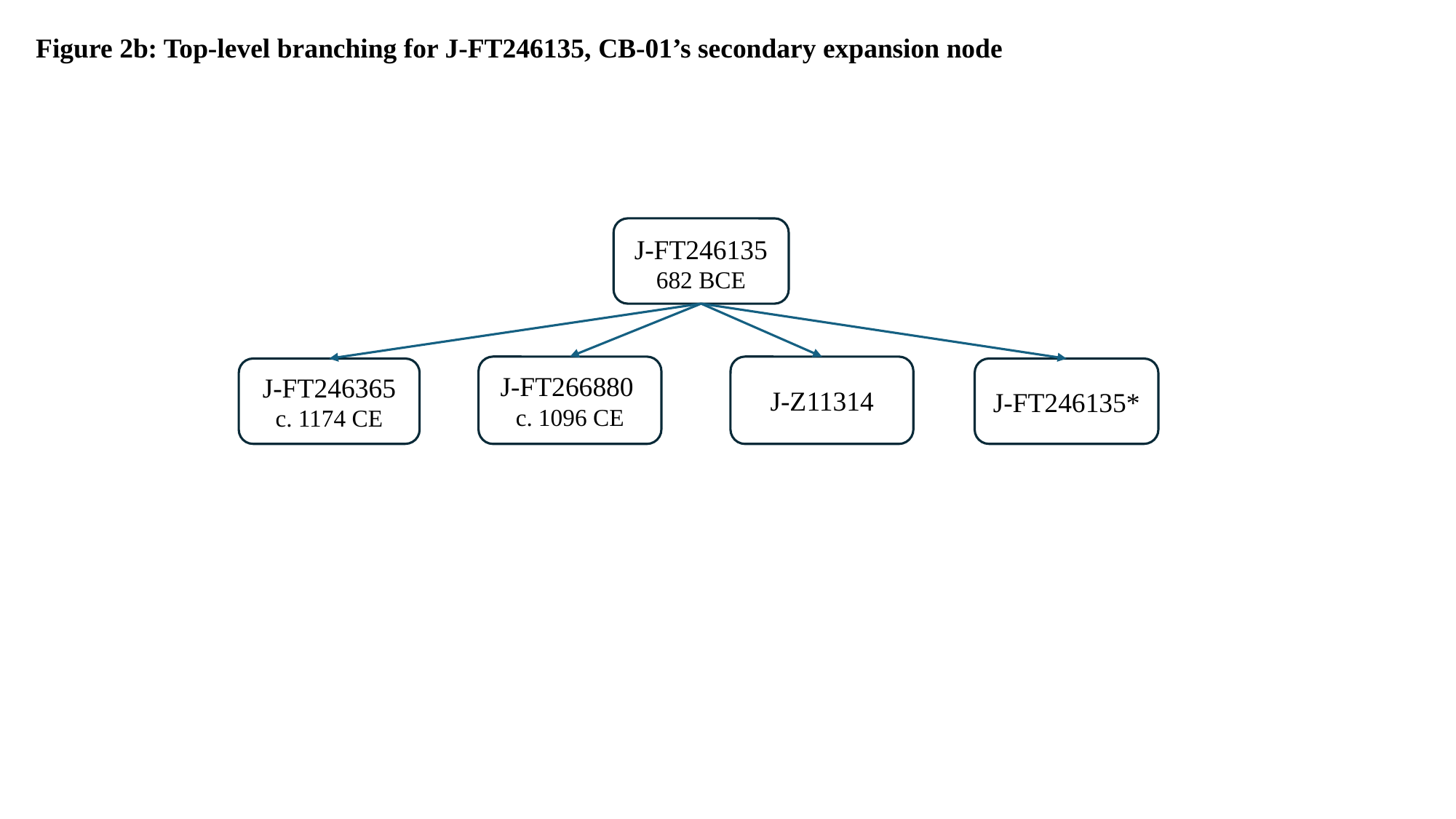

Figure 2b: Top-level branching for J-FT246135, CB-01’s secondary expansion node
J-FT246135
682 BCE
J-FT266880
c. 1096 CE
J-Z11314
J-FT246365
c. 1174 CE
J-FT246135*

### Slide 4
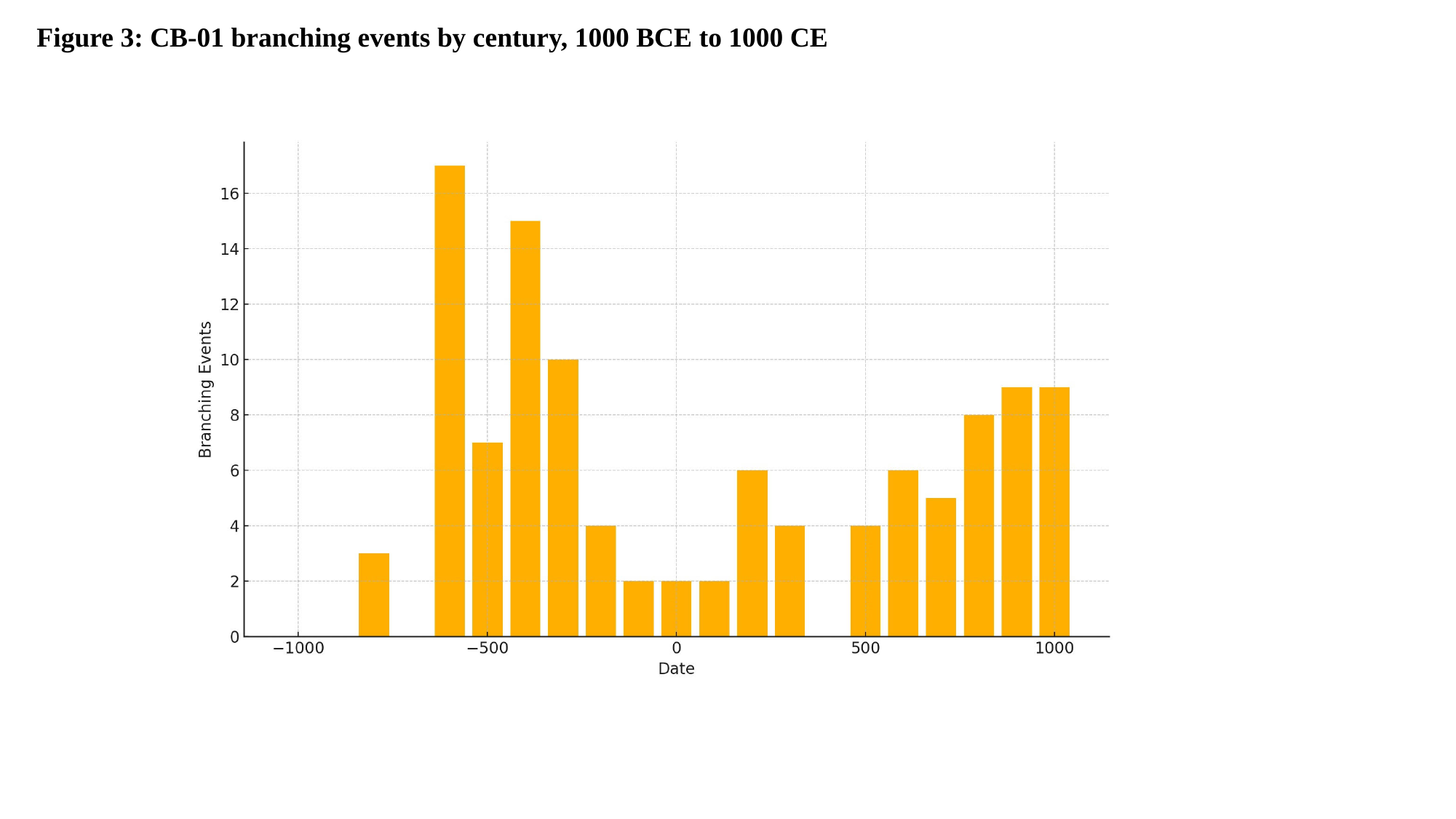

Figure 3: CB-01 branching events by century, 1000 BCE to 1000 CE

### Slide 5
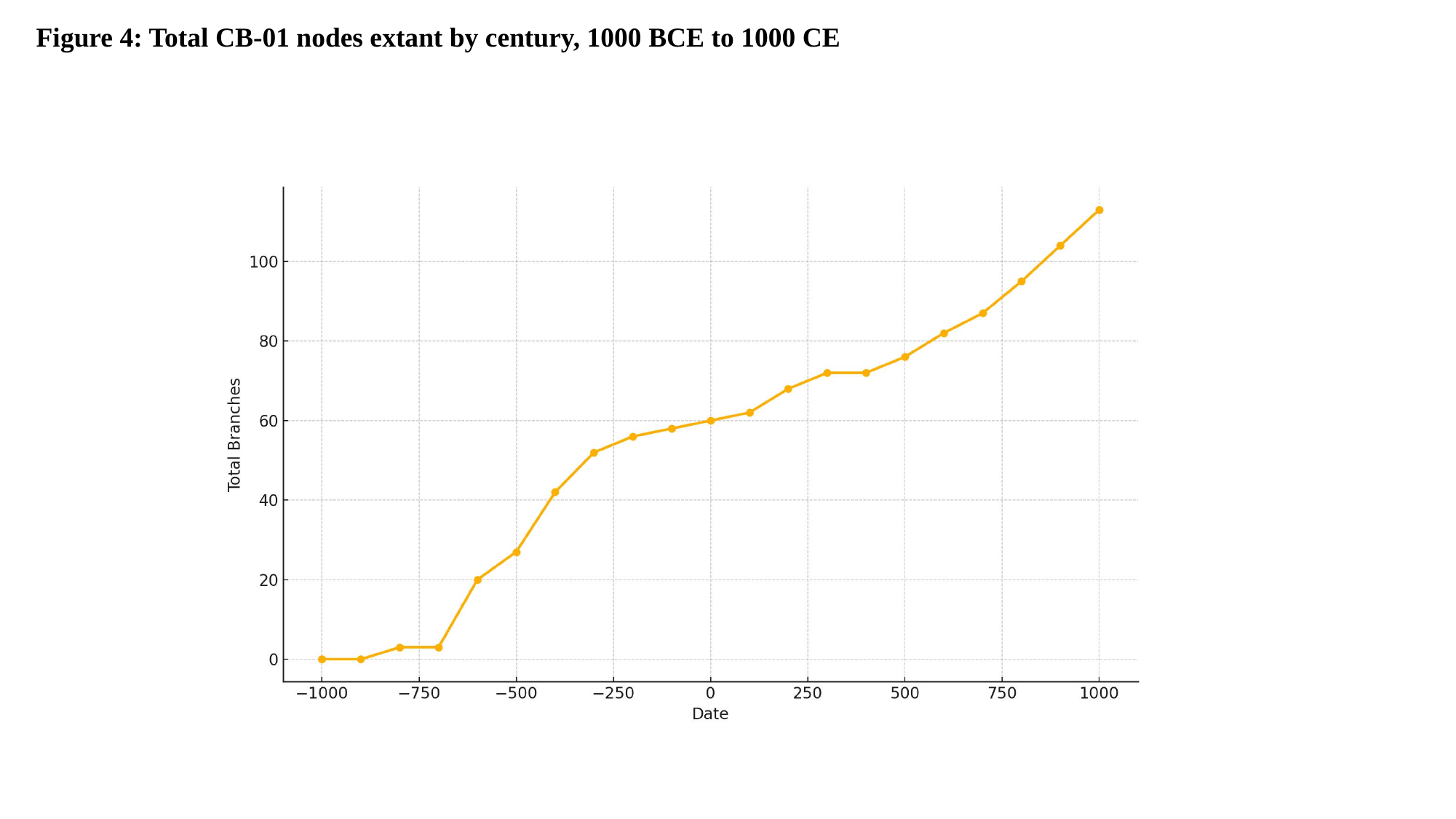

Figure 4: Total CB-01 nodes extant by century, 1000 BCE to 1000 CE

### Slide 6
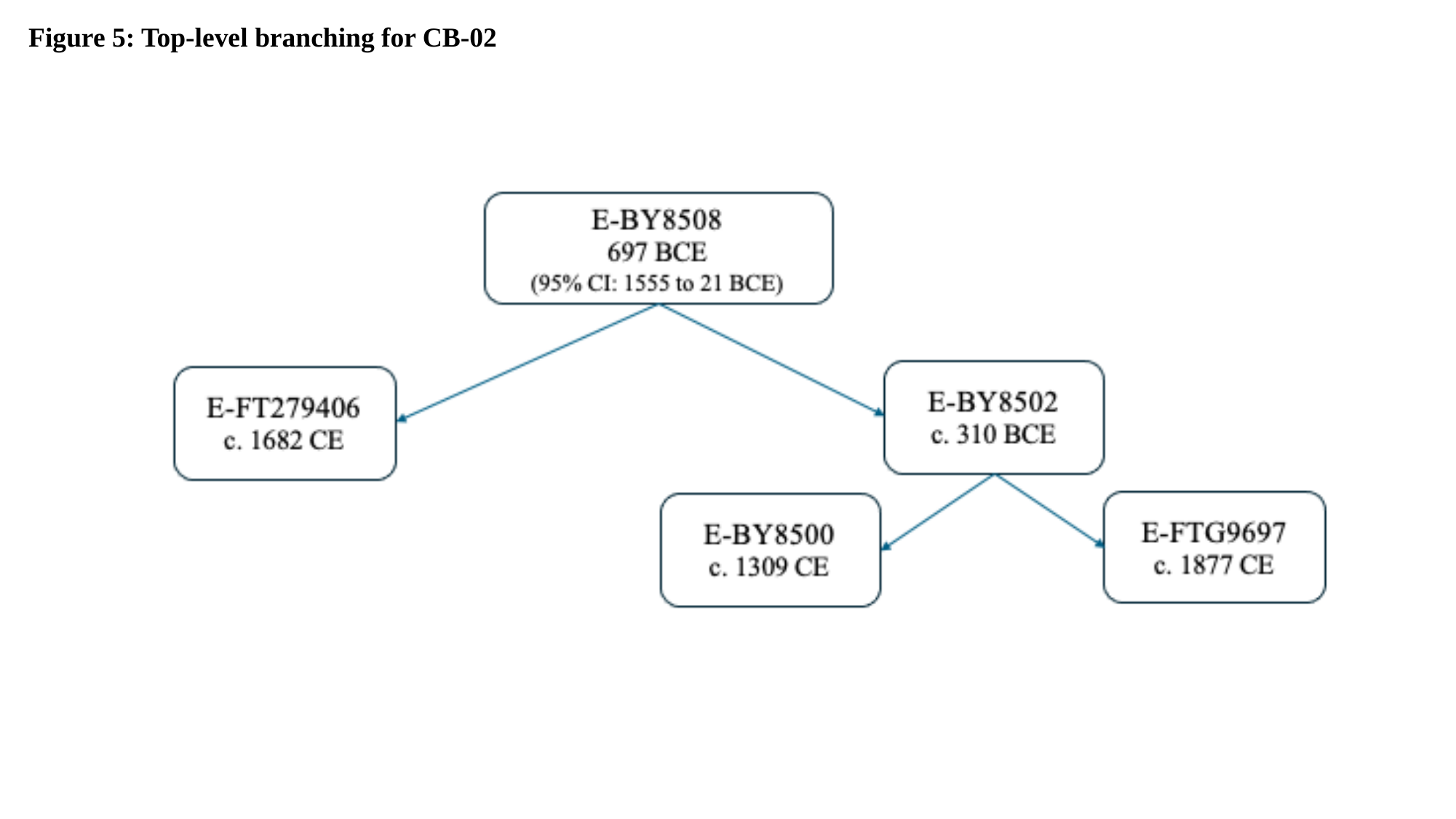

Figure 5: Top-level branching for CB-02

### Slide 7
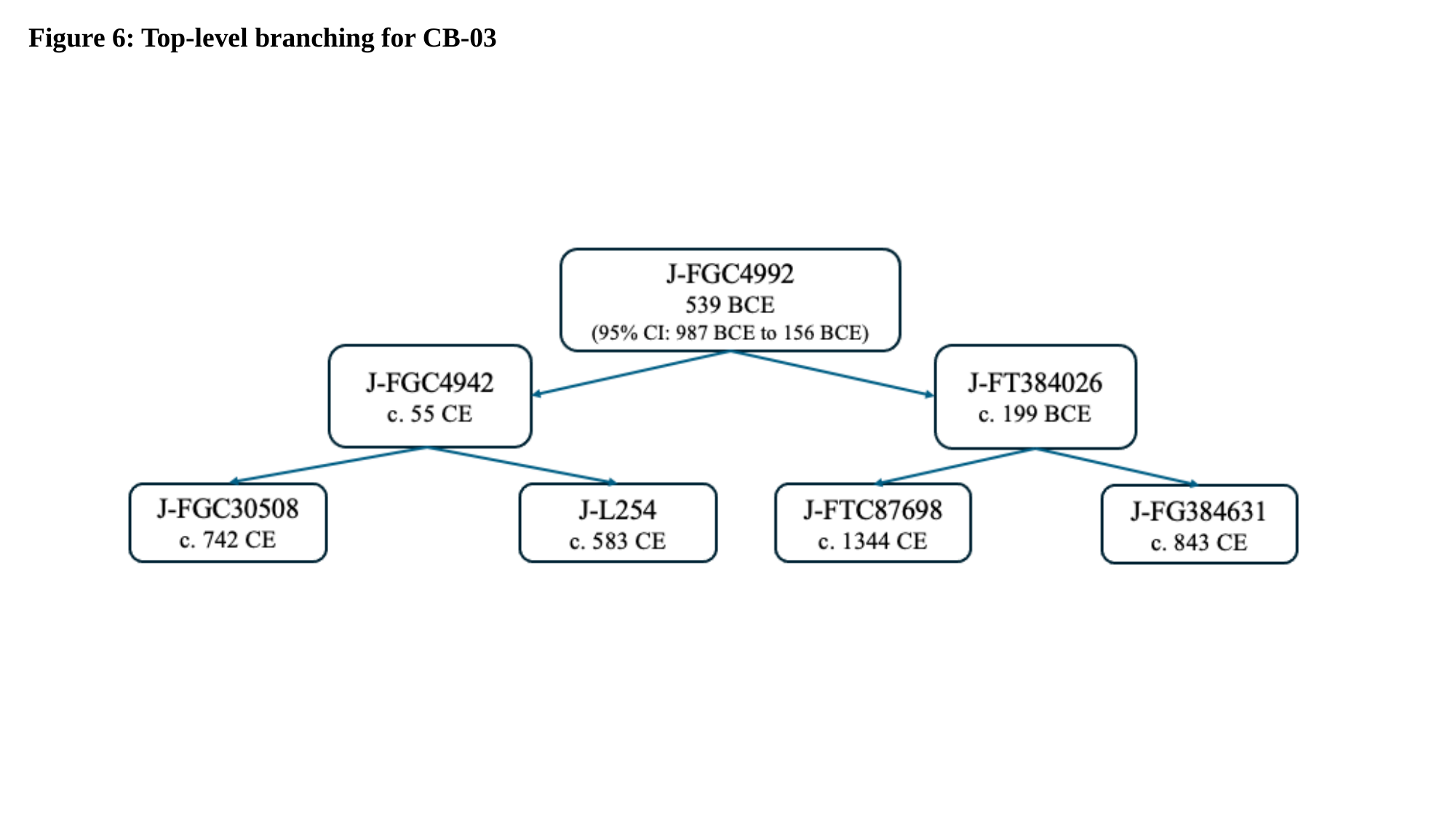

Figure 6: Top-level branching for CB-03

### Slide 8
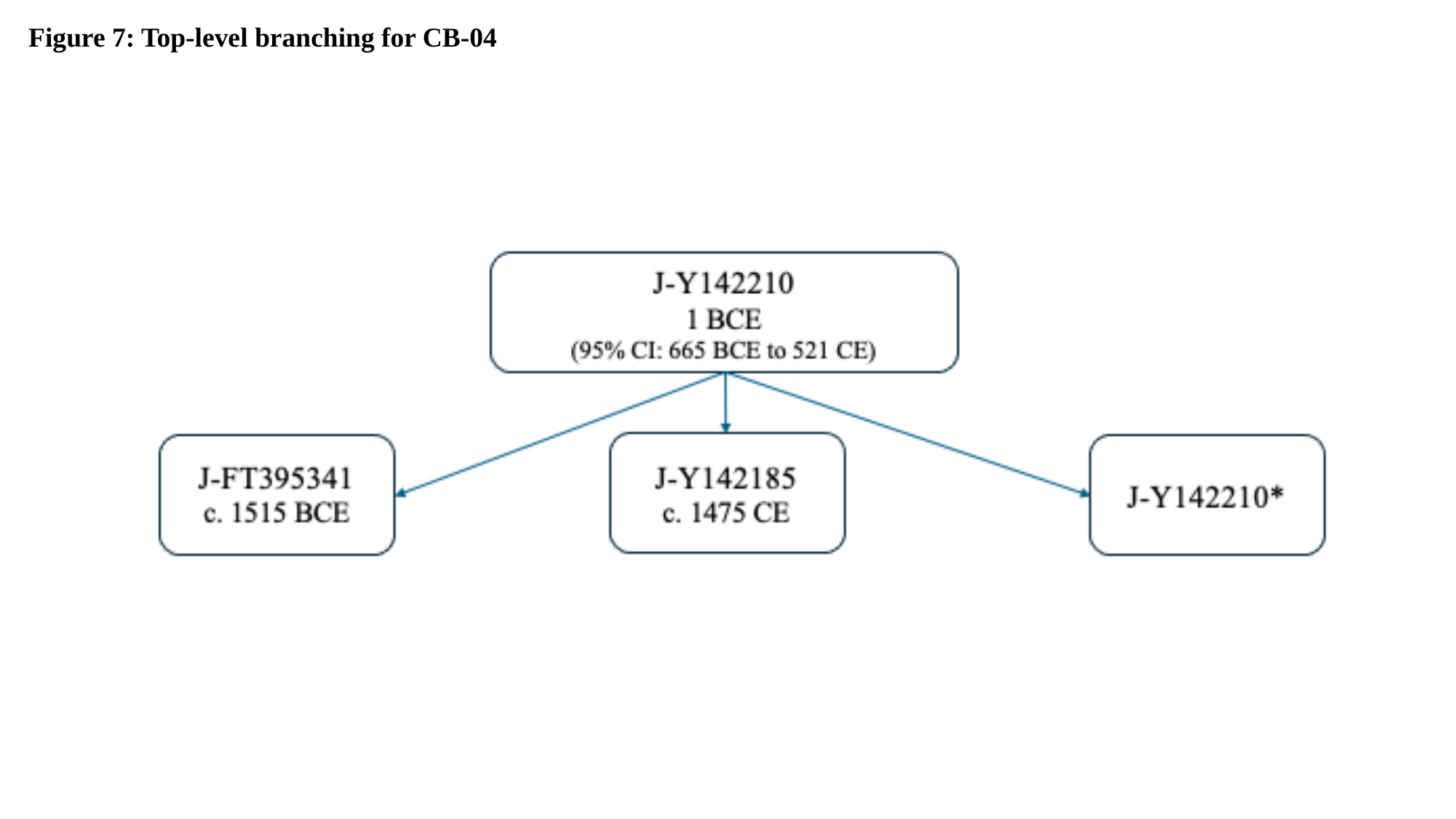

Figure 7: Top-level branching for CB-04

### Slide 9
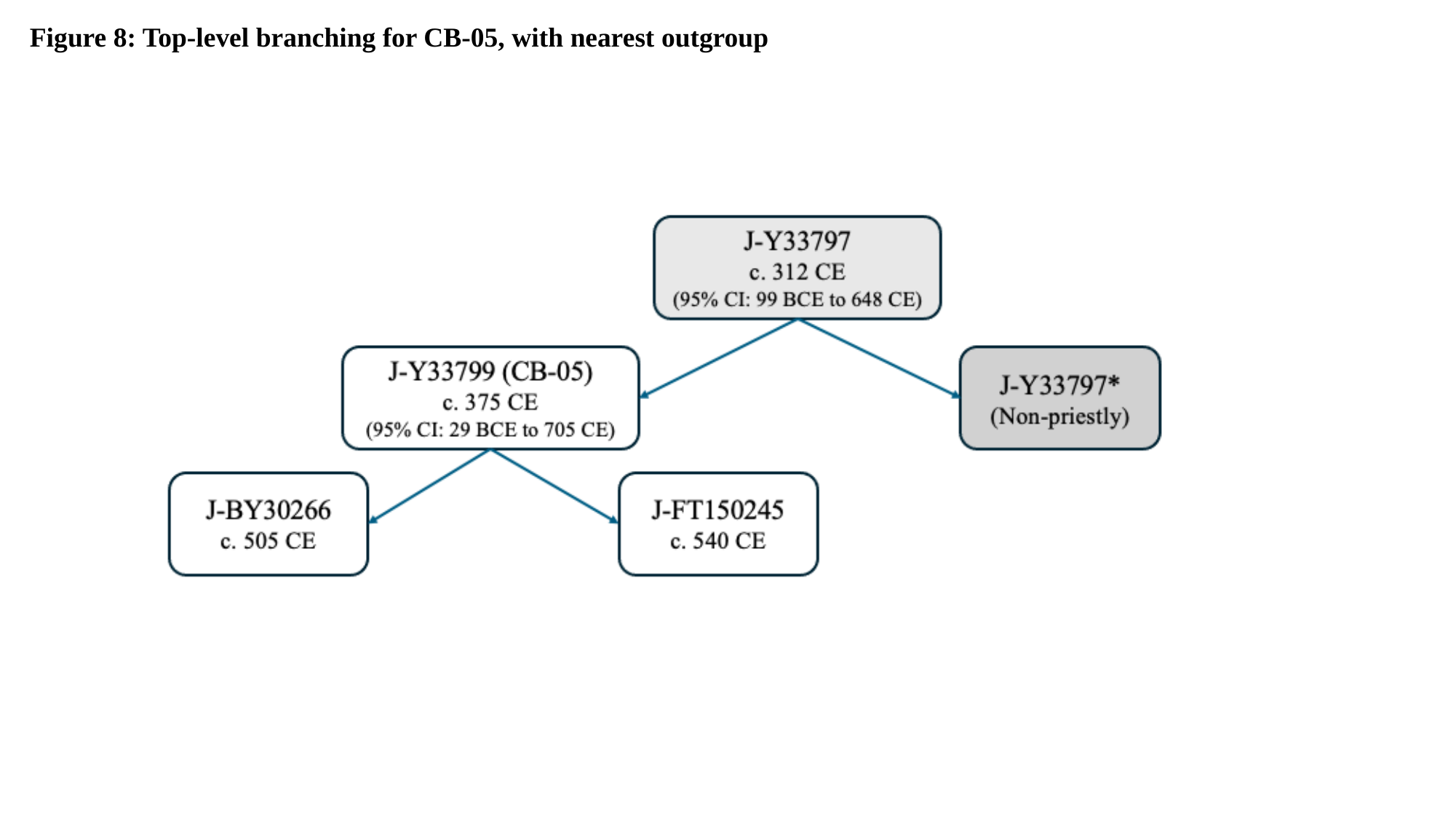

Figure 8: Top-level branching for CB-05, with nearest outgroup

### Slide 10
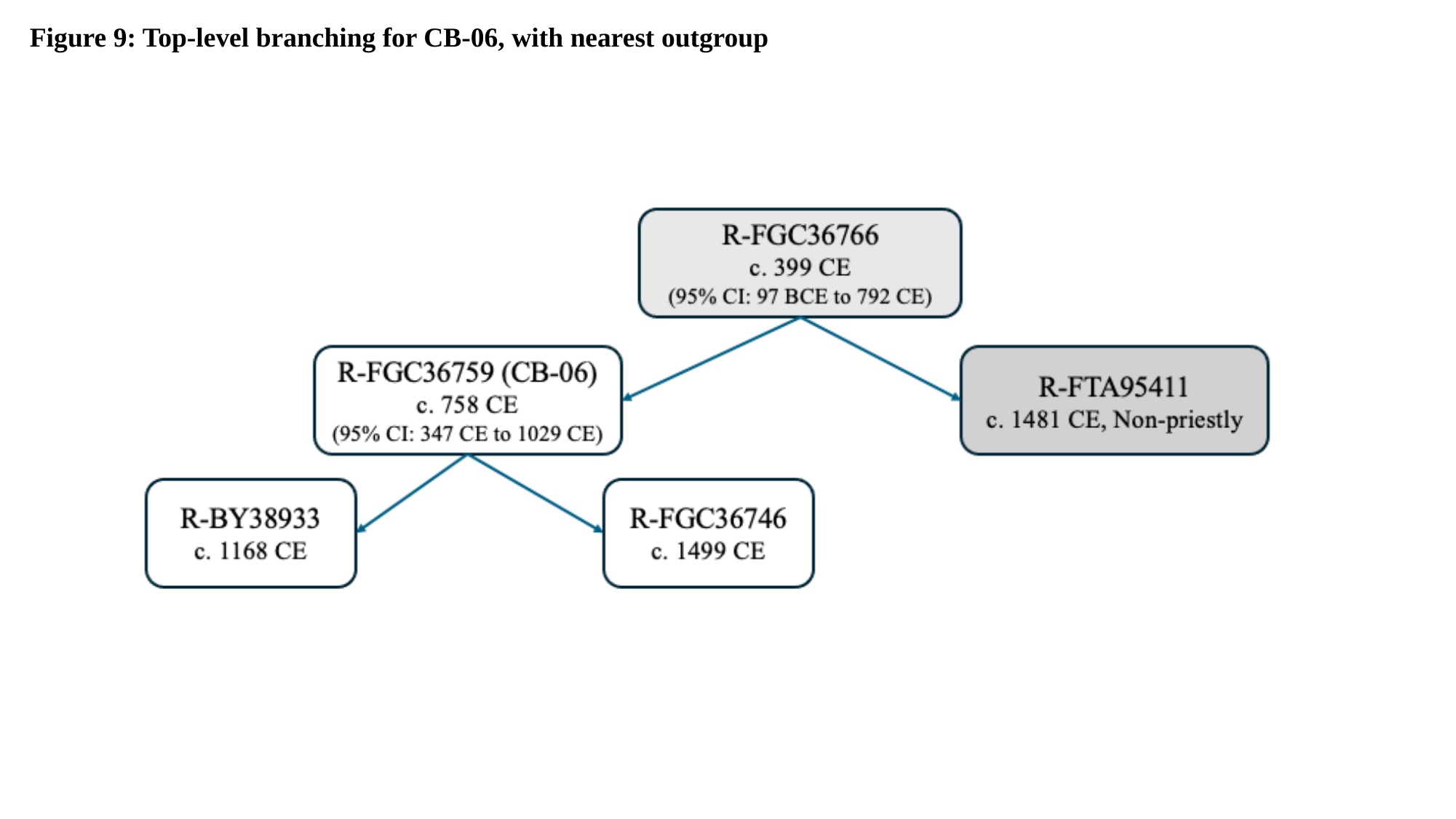

Figure 9: Top-level branching for CB-06, with nearest outgroup

### Slide 11
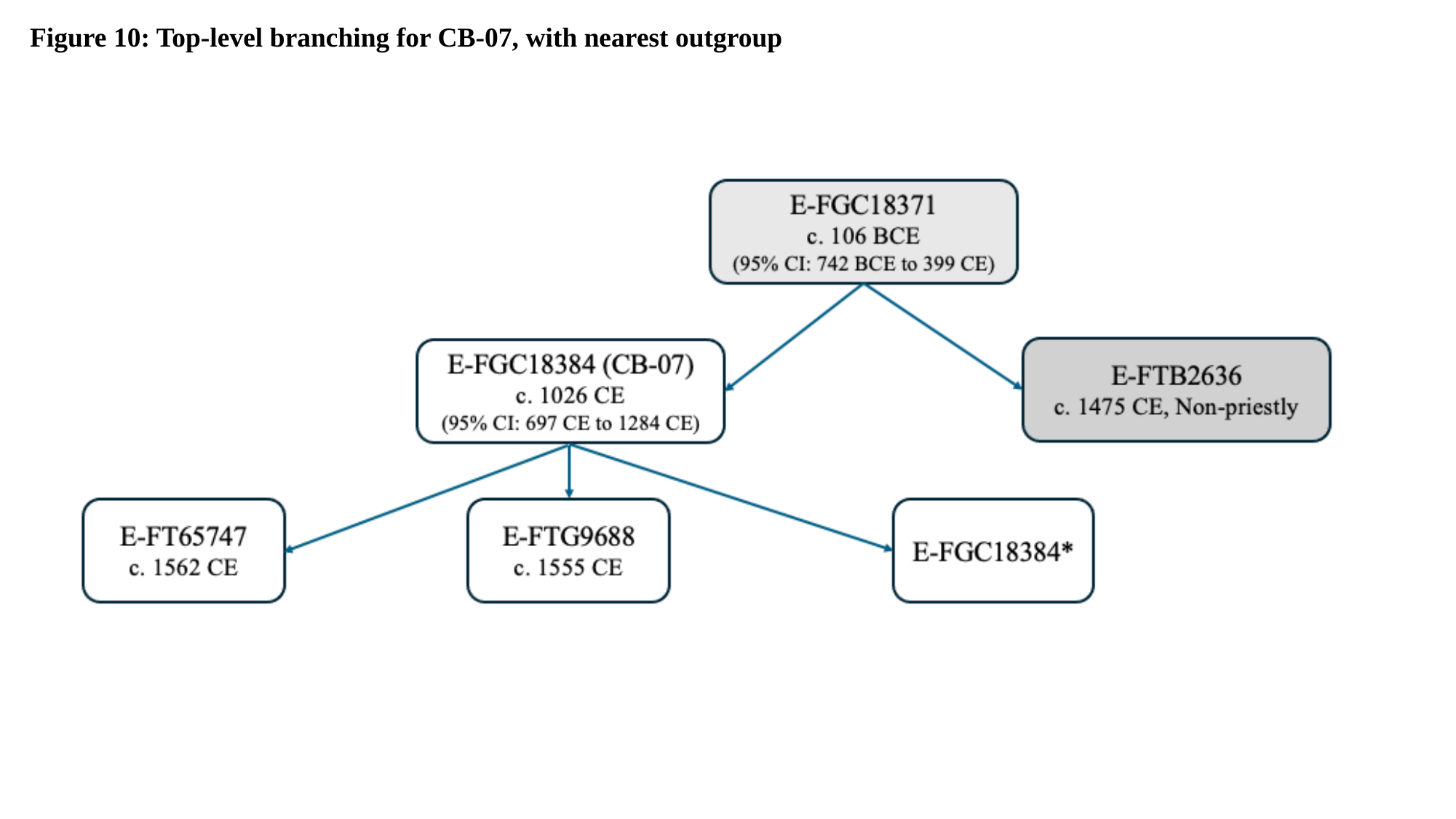

Figure 10: Top-level branching for CB-07, with nearest outgroup

### Slide 12
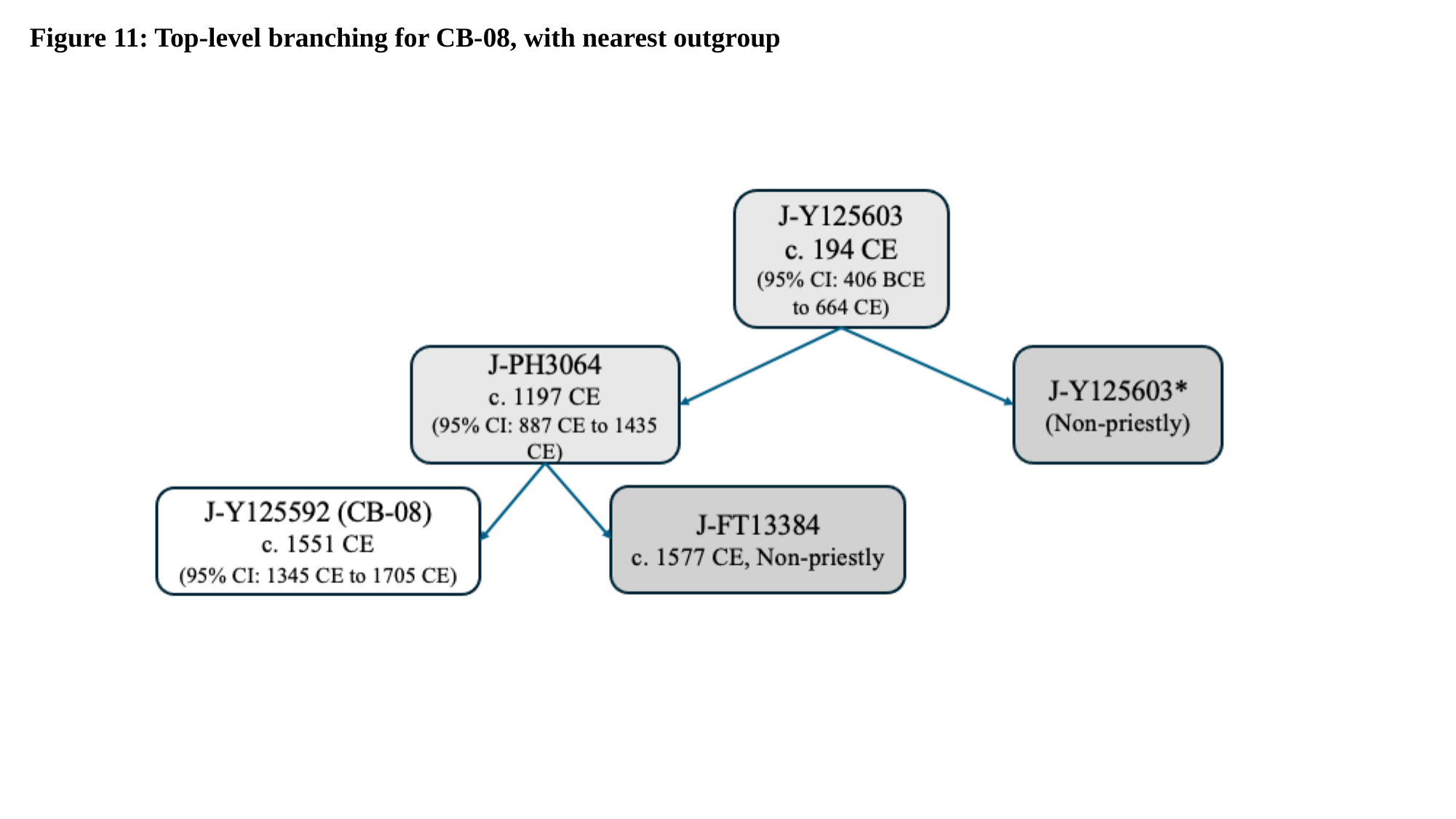

Figure 11: Top-level branching for CB-08, with nearest outgroup

### Slide 13
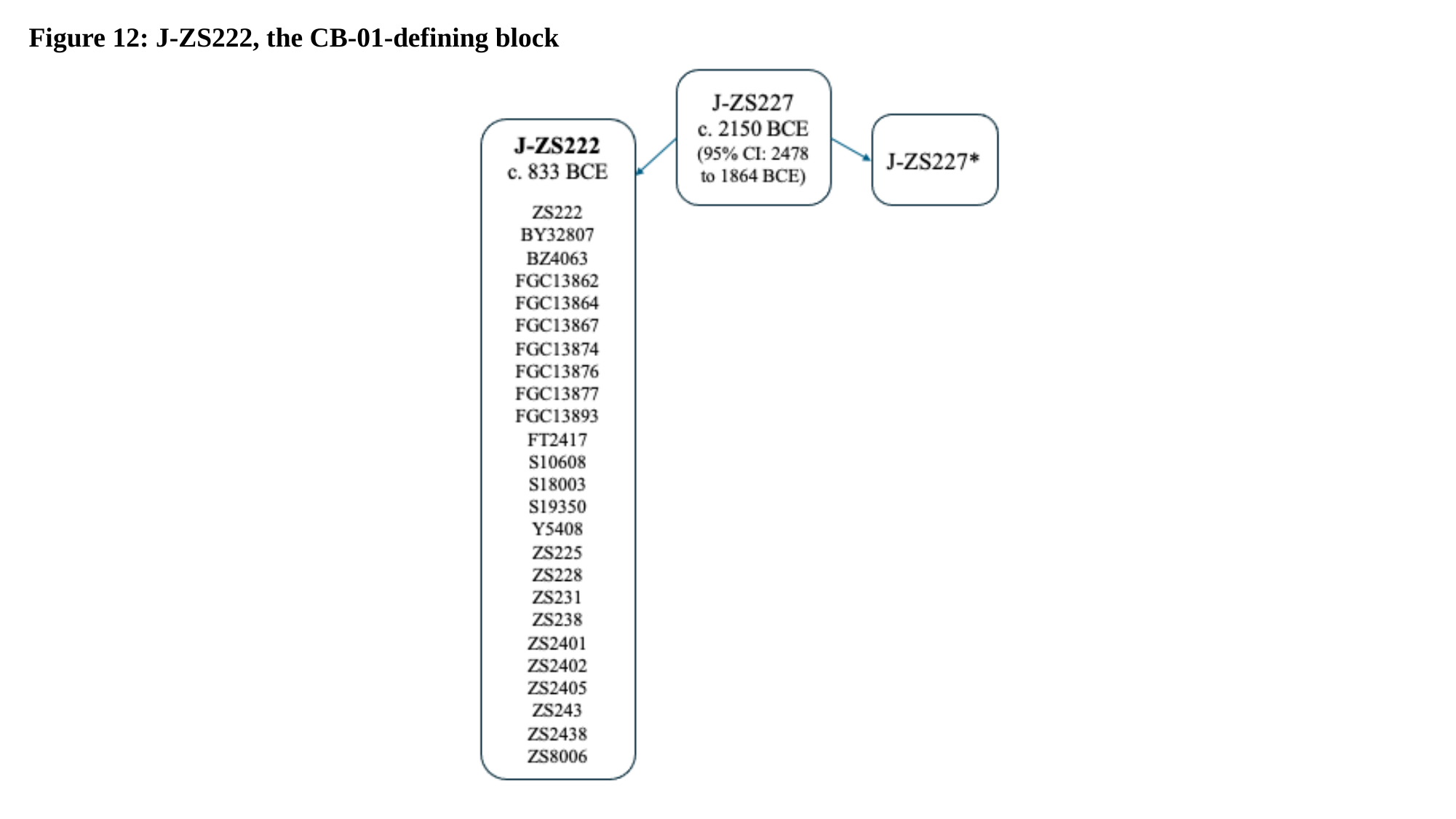

Figure 12: J-ZS222, the CB-01-defining block
