## Supplementary material for "Revisiting the lineages of the Cohanim using data from next-generation sequencing": Figure Legends

Figure 1 depicts CB-01's (J-ZS222) split into three primary branches, which share a common ancestor c. 833 BCE (95% CI: 1289 to 440 BCE). Two of these branches are named—J-Z18271 and J-FT246134, both coalescing to common ancestors c. the 7th century BCE—and one is represented by a single basal individual (J-ZS222*).

Figure 2a depicts the top-level branching for CB-01's largest primary branch, J-Z18271, into 11 primary sub-branches of its own. Estimated coalescence dates for each of these 11 branches is given; all are derived from FTDNA, with the exception of J-Y191714, for which only YFull generates a TMRCA estimate.

Figure 2b depicts the top-level branching for CB-01's second-largest primary branch, J-FT246135, into 11 primary sub-branches of its own. Estimated coalescence dates for these sub-branches is given when possible; all are derived from FTDNA.

Figure 3 depicts the number of distinct branching events within CB-01, beginning with its initial branching event in the 9th century BCE, and continuing through 1000 CE. Branching events following this medieval date are not included in the present chart. A peak in branching is identified between 700 and 300 BCE, followed by a slowing of branching, followed by a steady resumption of branching events beginning in late antiquity. This analysis is necessarily limited to branching events involving tested modern individuals.

Figure 4 depicts the number of nodes of CB-01, estimated to have been extant by the end of each century, beginning with its initial coalescence in the 9th century BCE, and continuing through 1000 CE. Growth following this medieval date is not included in the present chart. Once again, a the number of extant nodes grows quickly between 700 and 300 BCE, followed by a slowing of branching, followed by a steady increase in growth beginning in late antiquity. This analysis is necessarily limited to branching events involving tested modern individuals.

Figure 5 depicts CB-02's (E-BY8508) divergence into two primary branches, which share a common ancestor c. 697 BCE (95% CI: 1555 to 21 BCE). For the older of these two branches (E-BY8502), further branching is shown.

Figure 6 depicts CB-03's (J-FGC4992) divergence into two primary branches, which share a common ancestor c. 539 BCE (95% CI: 987 to 156 BCE). Both branches—J-FGC4942 and J-FT384026, coalescing to common ancestors in the classical period. Further historically relevant sub-branching is shown for both primary branches.

Figure 7 depicts CB-04's (J-Y142210) split into three primary branches, which share a common ancestor c. 1 BCE (95% CI: 665 BCE to 521 CE). Two of these branches are named—J-FT395341 and J-Y142185, both coalescing to common ancestors c. 15th-16th centuries CE—and one is represented by a single basal individual (J-Y142210*).

Figure 8 depicts CB-05's (J-Y33799) internal structure, and includes its closest Jewish, but non-Cohen-affiliated, outgroup.

Figure 9 depicts CB-06's (R-FGC36759) internal structure, and includes its closest Jewish, but non-Cohen-affiliated, outgroup.

Figure 10 depicts CB-07's (E-FGC18384) split into three primary branches, which share a common ancestor c. 1026 CE (95% CI: 697 to 1284 CE). Two of these branches are named—E-FT65747 and E-FTG9688, both coalescing to common ancestors c. 15$^{th}$-16$^{th}$ centuries CE—and one is represented by a single basal individual (E-FGC18384*). CB-07's closest Jewish, but non-Cohen-affiliated outgroup, is also shown.

Figure 11 depicts CB-08's (J-Y125592) position within a wider Jewish branch (J-Y125603), whose other distinct sub-branches lack Cohen affiliation.

Figure 12 depicts CB-01 (J-ZS222) and the block of 25 SNPs that define it, relative to the upstream layer, J-ZS227. CB-01's divergence from a basal J-ZS227* individual is estimated to date to c. 2150 BCE (95% CI: 2478 to 1864 BCE).
