## Supplementary Informaton for "Revisiting the lineages of the Cohanim using data from next-generation sequencing"

**Supplementary Information**

*Downstream structure of J-Z18271, the largest sub-branch of CB-01*

**J-Y191714** (c. 1725 CE) is defined by two Pontic Greek, non-Jewish individuals, with matches to many other Pontic Greek individuals, some of whom have a tradition of being “priests”.

**J-FT280381** (c. 1300 CE) is defined by three Yemenite Cohanim.

**J-FT303739** (c. 850 CE) is defined by a Bulgarian Jew, a Greek Jew, and a Turkish Jew, some of whom have confirmed traditions of priestly origin.

**J-ZS237** (c. 100 CE) is defined by a Peruvian individual and several Southern Italian individuals, none of whom are known to have recent Jewish ancestry.

**J-ZS2458** (c. 100 CE) includes Iraqi, Mountain, Afghan, and Crimean Karaite Cohanim.

**J-BY64521** (c. 400 BCE) includes Iraqi, Iranian, Syrian, and Bukharian Cohanim, as well as Lebanese and Palestinian Christians.

**J-Y146901** (c. 500 BCE) is defined by an Ashkenazi Jew and a Greek Sephardic Jew confirmed to have traditions of priestly origin, as well as individuals of various Hispanic and Iberian origins, an English individual, and a Turkish individual.

**J-S17446** (c. 500 BCE) is defined by Ashkenazi, Moroccan, Algerian, and Yemenite Cohanim, as well as multiple clusters of Hispanic-origin individuals. This sub-branch includes the largest Ashkenazi cluster of CB-01, under J-S12192.

**J-ZS2434** (c. 500 BCE) is defined by Syrian and Yemenite Jews of unknown priestly status and a cluster of Ashkenazi non-Cohanim, and also includes a Sicilian individual with no known recent Jewish ancestry.

**J-FT34605** (c. 550 BCE) is defined by a Turkish Sephardic Cohen, as well as several Hispanic individuals, a Kazakh, and a Circassian.

**J-FT157560** (c. 600 BCE) is the oldest and most structurally complex sub-branch of J-Z18271, and of CB-01 at large, spanning Western and Eastern Jewish Diaspora groups, and containing multiple distinct Ashkenazi, Moroccan, and Eastern Sephardic sub-branches/clusters. This sub-branch includes Ashkenazi, Moroccan, Algerian, Tunisian, Greek, Turkish, Syrian, and Iraqi Cohanim, as well as individuals of various Hispanic and Iberian origins.

The convergence of all of these aforementioned currently Jewish sub-branches, all but one of them associated with Cohen status, back to c. 700 BCE, suggests that the common ancestor of J-Z18271 was likely to have been a Cohen.

*Internal structure and phylogenetic affinities of CB-02 to CB-09*

**CB-02**’s nearest upstream connection is to a branch containing two Armenian individuals, estimated to date to approximately 1450 BCE. One level above, currently defined as E-CTS8411, CB-02 branches off around 1600 BCE, linking to other distinctly Jewish Y-DNA branches as well as to modern Armenian, Hispanic, and English lineages.

E-FT279406, a young branch dating to c. 1682 CE, has only been observed in Bukharian Jews from Uzbekistan with a priestly tradition. E-BY8502, by contrast, splits c. 310 BCE into two sub-branches: E-FTG9697 (c. 1877 CE), a genealogically recent lineage found in Yemenite Jews with Cohen traditions, and E-BY8500 (c. 1309 CE), which is present among Syrian, Turkish, and Egyptian Jews with priestly heritage, as well as among some Hispanic individuals with no present tradition of Jewish descent.

Given that all known branches of CB-02, converging back to c. 697 BCE, are associated with priestly status, E-BY8508 may have been associated with priestly status in classical Judaea (c. late 1^st^ millennium BCE to 70 CE), potentially even as early as the monarchic period in ancient Israel and Judah (c. 10th century BCE to 586 BCE).

**CB-03**’s closest upstream connection dates back to the Neolithic period (c. 6000 BCE) at the J-PF5366 phylogenetic level, which includes a variety of European, Near Eastern, and Caucasian individuals, along with an individual from Roman-era Anatolia.

CB-03 (c. 539 BCE) contains two primary sub-branches: J-FGC4942 and J-FT384026. CB-03 (c. 55 CE) is found in Ashkenazi, Moroccan, Greek, and Turkish Jews, most of whom identify as Cohanim (**Figure 6**). J-FT384026 (c. 199 BCE) has been identified in two Mountain (Caucasian) Jewish individuals and two Cochin (Indian) Jewish individuals. It must be noted, in this case, that the priestly status of the two Cochin Jewish branch members is unknown, and only one of the two Mountain Jews has a priestly tradition. While this requires us to exercise caution in including J-FT384026 under the scope of CB-03, we also note that this picture may be affected by the extreme rarity of priestly tradition among Jewish communities in the North Caucasus. While we conjecture that the lower rate of Cohen identification within this sub-branch reflects partial loss of Cohen status, it may also represent a later, partial acquisition of Cohen status by one of its Mountain Jewish descendants.

The convergence of J-FGC4992’s Western and Eastern Jewish sub-branches around 539 BCE, along with its clear Near Eastern phylogenetic origins, suggests a likely presence in ancient Judaea. While a priestly identification is reasonably likely for all of J-FGC4992, and thus we define it as CB-03, we apply this broader definition with some caution, and couple it with a more restricted, higher-confidence possible definition of CB-03 as the younger (c. 55 CE), Western Jewish J-FGC4942 sub-branch.

**CB-04**’s closest upstream branch, J-FT284582, connects it to an Armenian individual around approximately 750 BCE. One levels further up, CB-04 is connected to various Near Eastern lineages c. 2050 BCE.

The majority of tested individuals within CB-04 fall within one young sub-branch, J-Y142185, which is estimated to have a common ancestor c. 1475 CE; all of these individuals report Afghan or Iranian Jewish origin and priestly tradition. This sub-branch connects c. 1 BCE with a single young branch containing an Iraqi Jewish Cohen and an Armenian individual, as well as with a basal branch defined by a single Iraqi Kurdish Jewish individual, whose priestly status is uncertain (**Figure 7**).

CB-04’s age suggests a common Jewish ancestor in classical Judaea, and a priestly tradition dating back at least to classical times. However, given the small sample size outside of the J-Y142185 sub-branch, and the uncertain Cohen status of the basal Iraqi Kurdish Jewish individual, this assignment must be made with some caution.

**CB-05** shares the J-Y33797 level with a Moroccan Jewish individual without a priestly tradition, a connection estimated to approximately 300 CE (**Figure 8**). One level more remotely, J-Y33797 converges with a Swiss and Hungarian lineage, a connection dating to approximately 1200 BCE. Further upstream, at J-Z34474, CB-04 diverges around 1400 BCE from branches including Italian, German, Portuguese, and Anglo-American individuals, as well as an Etruscan individual from Iron Age Italy [30].

Currently, CB-05 diverges into two exclusively Ashkenazi Jewish sub-branches, with the majority of individuals in both identifying as Cohanim, c. 375 CE (half a millennium prior to the earliest attested Ashkenazi communities). The association of J-Y33799 with priestly status suggests that this branch may have acquired Cohen status in the late classical Western Jewish Diaspora, namely, somewhat later than its earliest attested Jewish status. Its phylogenetic context and relatively close European connections suggest antique European introgression.

**CB-06**’s closest upstream branch (defined by the R-FGC36766 block and its 12 associated SNPs) includes several Syrian Jews without Cohen affiliation, a connection which dates to around 400 CE (**Figure 9**). At the R-L944 level, CB-06 connects around 450 BCE to Lebanese, Turkish, and German-American lineages.

CB-06 splits c. 758 CE into two branches, both exclusively comprising Ashkenazi Jews, most of whom identify as Cohanim. The divergence from non-Cohen Syrian Jews c. 400 CE suggests that while the branch’s Jewish origins may date to classical times, its priestly pedigree may be more recent, dating to the late antique or early medieval period.

**CB-07**’s closest connection, defined by the SNP E-FGC18371 and its associated block of 9 SNPs, is to a small branch of Ashkenazi Jews without traditions of priestly descent, and dates to around 100 BCE (**Figure 10**). At the E-FGC18422 level, CB-07 connects around 1000 BCE to other Near Eastern and European branches. Moreover, a recently published sample from an Iron Age individual in southeastern Turkey/upper Mesopotamia was found to belong to the broader upstream branch E-FGC18401, thus sharing a common ancestor with CB-07 no later than 1350 BCE [31].

CB-07 diverges into three primary sub-branches c. 1026 CE, with one basal Iranian Jewish-defined branch, an Iraqi Jewish branch, and an Iranian-Afghan Jews in the other. The divergence from non-priestly Jews around 100 BCE suggests a likely common ancient Jewish origin, but does not resolve whether the common ancestor was of priestly status. The divergence from non-Cohen Ashkenazi Jews around 100 BCE suggests that while the branch’s Jewish origins may date to classical times or earlier, its priestly pedigree cannot be confidently traced beyond the medieval period.

**CB-08** is situated within a wider Jewish branch (J-Y125603) dating to around 200 CE, with members among Djerban, Tunisian, and Libyan Jews. However, the priestly status of these individuals is unclear in some cases, and negative in others. The branch’s next closest upstream connection is to a branch containing a Mexican individual, estimated to date to around 150 CE. Additionally, one level above (currently defined as J-M318), CB-08 connects around 750 BCE to a branch containing ethnically Italian and German individuals.

Given that priestly status can only be clearly associated with Djerban Jews at the J-Y125592 level, the priestly pedigree of CB-08 cannot be securely dated earlier than c. 1551 CE (**Figure 11**). However, the identification of several additional Tunisian and Libyan Cohanim within the J-M318 branch reported in Hammer et al. (2009) raises the possibility that CB-08’s priestly status might antedate the recent J-Y125592 level; however, this cannot be demonstrated definitively. Given its close upstream associations in Mediterranean Europe, it is possible that this branch is ultimately a marker of classical Mediterranean introgression in Libyo-Tunisian Jews, with possible subsequent acquisition of priestly status in the Diaspora.

**CB-09**’s closest upstream branch (E-BY188966) includes individuals of British, Central European, and Iranian descent dating to around 700 BCE. These European associations suggest potential classical Mediterranean introgression in Syrian or Mediterranean Jewish communities, but cannot establish the branch’s priestly association prior to late medieval/early modern times.
